# β-Coronavirus Nsp6 hijacks host ER translocation machineries into viral replication centers

**DOI:** 10.64898/2025.12.16.694738

**Authors:** Kari H. Ecklund, Robert G. Abrisch, Stanley Perlman, Gia K. Voeltz

## Abstract

β-coronaviruses evade host immune detection by replicating their genomes within double membrane vesicles (DMVs) derived from endoplasmic reticulum (ER) membranes. For example, SARS-CoV and CoV-2 encode three non-structural membrane proteins (Nsp 3, 4, and 6) which can remodel the ER to form DMVs. Here we test whether Nsps also function to recruit key host machineries required for viral replication and assembly within ER-derived DMVs. We use mouse hepatitis virus to study whether β-coronavirus Nsps coordinate ER remodeling with host machinery recruitment. We demonstrate that Nsp6 generates Nsp6-remodeled ER domains that sequester host ER insertases including the Sec61 translocon, EMC, and GEL complexes. FRAP and FLIP experiments confirm that Nsp6 domains remain continuous with the ER and do not restrict membrane protein diffusion, except for those insertases that are sequestered there by Nsp6. Together, these data demonstrate a dual role for Nsp6 in remodeling ER membranes and sequestering host translocation machinery away from the general ER and into DMVs.

## INTRODUCTION

β-coronaviruses assemble and replicate within double membrane vesicles (DMVs) made from remodeled ER membranes^1–4^. Electron microscopy (EM) of infected cells has revealed that DMVs are composed of zippered membranes and double membrane vesicles (DMVs that appear continuous with the ER and are often associated with ribosomes)^2,3,5^. Upon entering the cell, replicase genes are directly translated into two polyproteins, PP1a and PP1ab, which are cleaved into 16 non-structural proteins (Nsp 1-16)^6–8^. The viral transmembrane proteins Nsp3, Nsp4, and Nsp6 have been shown by both fluorescence and electron microscopy to be sufficient to remodel the ER membrane^9,10^. Ectopic co-expression of Nsp3 and Nsp4 is sufficient to induce DMVs in cell culture^9,11–13^. The additional co-expression of Nsp6 with Nsp3 and Nsp4 generates networks of clustered DMVs and zippered membranes reminiscent of viral infection^9,10^. These membrane rearrangements are proposed to function together, creating a barrier that could further restrict host factors from accessing DMVs^10^. This strategy allows β-coronaviruses to cloak themselves in the ER and hide from detection in the cytoplasm^2^. What remains unclear is by what mechanism do host machinery insertases gain access to ER-derived DMVs and translocate the viral nonstructural membrane proteins Nsp3, Nsp4, and Nsp6 for efficient DMV biogenesis and viral replication.

The Sec61 translocon is the canonical protein-conducting channel that is bound to active ribosomes to translocate nascent polypeptide chains into the ER and into the secretory pathway^14–18^. β-coronavirus replication is highly sensitive to inhibitors of the Sec61 translocon including Apratoxin S4 and Ipomoeassin-F^19–22^. This is not surprising given that the β-coronavirus genome encodes several TMD-containing proteins critical for both DMV biogenesis and virion assembly but it reveals that viral protein production requires at a minimum the Sec61 translocon. Here, we investigate how the β-coronavirus MHV rearranges ER membranes to form DMVs capable of supporting viral replication. We show that a single viral nonstructural membrane protein Nsp6 has dual and complementary functions during DMV biogenesis. It can hijack select host translocation machineries into Nsp6-remodeled ER domains that are continuous with the ER but from which they cannot escape.

## RESULTS

### MHV nonstructural proteins remodel ER domains in hepatocyte cells

Ectopic co-expression of three virally encoded non-structural membrane proteins Nsp3, Nsp4, and Nsp6 from either SARS-CoV or SARS-CoV-2 is sufficient to remodel the ER membrane into domains that morphologically resemble the DMVs formed following viral infection in HEK293T-ACE2 and HeLa cells^9,10^. These three proteins contain multiple transmembrane domains (TMDs) (see cartoon in Figure 1A) which target them to the ER where they induce dramatic rearrangement of host membranes to form DMVs. We tested whether ectopic expression of the corresponding three non-structural proteins (Nsp3, Nsp4, and/or Nsp6) encoded by mouse hepatitis virus (MHV, strain JHM^IA,^ herein referred to as MHV23 would similarly remodel the ER membranes in murine FL83B hepatocytes, a mouse cell line with a large, morphologically flat cytoplasm suitable for our cellular measurements. FL83B cells were transiently co-transfected with mNe-Nsp3 (green), mCh-Nsp4 (yellow), and HALO-Nsp6 (magenta, conjugated to JFX650) along with either an ER lumenal (BFP-KDEL, blue) or ER membrane (BFP-Sec61β, blue) protein to visualize viral non-structural membrane protein localization relative to two well-characterized general ER markers. Indeed, FL38B cells expressing Nsp3, Nsp4 and Nsp6 produced spherical domains (Figure 1B and C) resembling those observed upon expression of Nsp proteins in many cell types for MHV and other β-coronaviruses^9,10,24^. Notably, we observed localization of Nsp3/4 domains in which they appeared “wrapping” Nsp6 domains, consistent with a proposed function of Nsp6 in clustering and organizing DMVs^10^. These domains were devoid of the lumenal ER marker BFP-KDEL (Figure 1B) consistent with previous reports that lumenal proteins are excluded by Nsp-induced membrane zippering^10^. In stark contrast, Nsp3/4/6 domains were highly enriched for the ER membrane protein Sec61β (Figure 1C). These data show that ectopic expression of three MHV non-structural membrane proteins (Nsp3, Nsp4, and Nsp6) remodels ER membranes in murine hepatocytes and thus provides a model system to dissect the roles of individual Nsp proteins in DMV formation.

**Figure 1:**
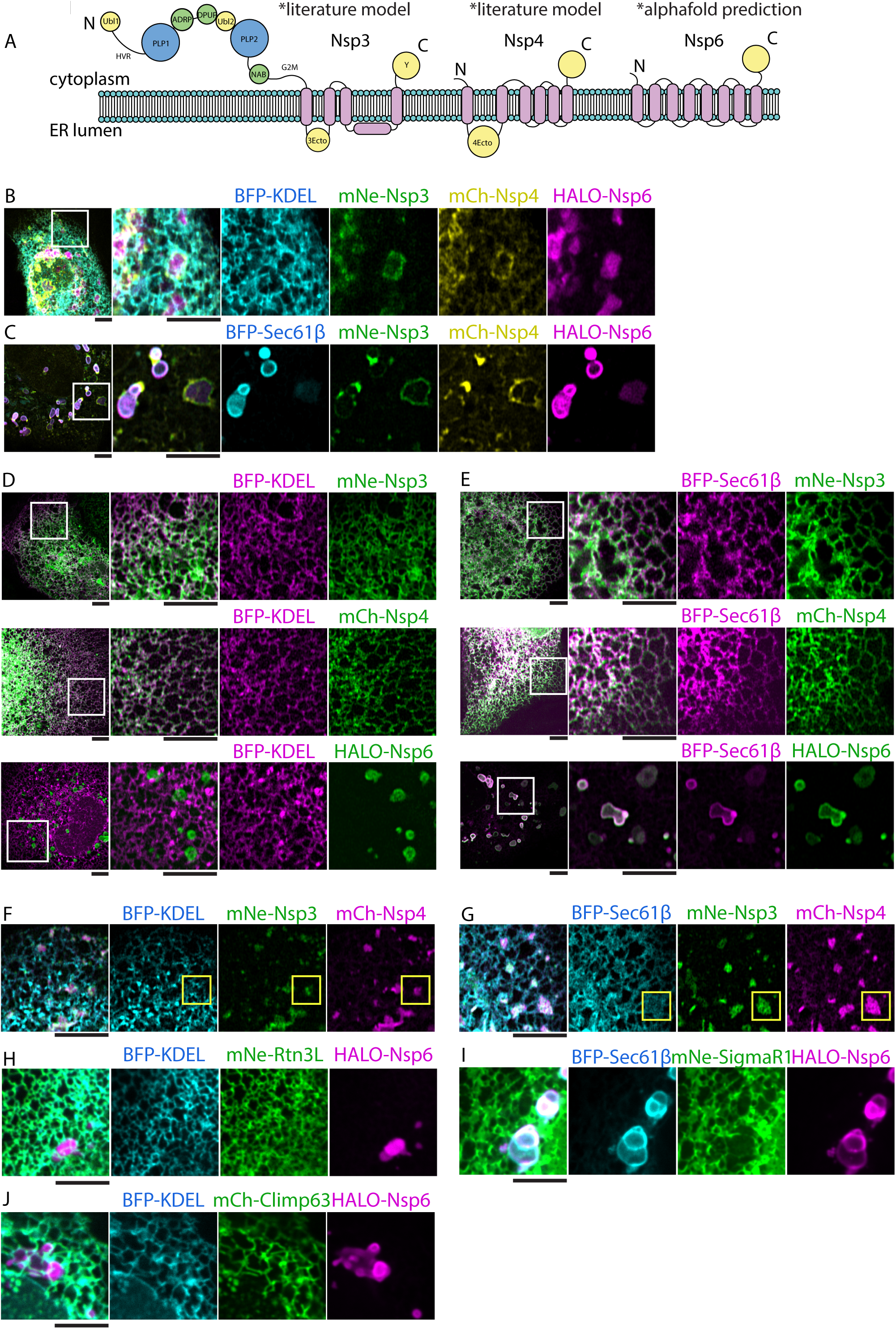
MHV nonstructural proteins remodel ER domains in hepatocyte cells. (A) (D) Cartoon of domain organization and topology of MHV nonstructural proteins Nsp3-6 derived from literature models and alphafold 3 predictions^90^. (B-C) Representative images of FL83B cells co-transfected with (B) mNe-Nsp3 (green), mCh-Nsp4 (yellow), HALO-Nsp6 (JFX650 conjugated, magenta) and with either (B) BFP-KDEL (blue) or (C) BFP-Sec61β (blue). (D) Images of FL83B cells co-transfected with BFP-KDEL (magenta) and mNe-Nsp3 (top, green), mCh-Nsp4 (middle, green), or HALO-Nsp6 (conjugated to JFX650, bottom, green). (E) Images of FL83B cells co-transfected with BFP-Sec61β (magenta) and mNe-Nsp3 (top, green), mCh-Nsp4 (middle, green), or HALO-Nsp6 (conjugated to JFX650, bottom, green). (F-G) Images of FL83B cells co-transfected with BFP-KDEL (F, blue) or BFP-Sec61β (G, blue) and with mNe-Nsp3 (green) and mCh-Nsp4 (magenta). (H-J) Images of FL83B cells co-transfected with BFP-KDEL (H and J, blue) or BFP-Sec61β (I, blue) and Halo-Nsp6 (magenta) with either (H) mNe-Rtn3L, (I) mNe-SigmaR1, or (J) mCh-Climp63 (green). Inset white boxes shown in left panels are magnified in panels on the right. Scale bars = 5 µm.

Next, we tested whether the individually expressed MHV Nsp proteins are sufficient to remodel ER membranes. FL83B cells were co-transfected with either mNe-Nsp3, mCh-Nsp4, or HALO-Nsp6 (each shown in green) along with either the lumenal ER marker BFP-KDEL or BFP-Sec61β (shown in blue) and were imaged live at 24 hours post transfection. Expression of mNe-Nsp3 or mCh-Nsp4 alone did not alter ER structure or promote the formation of specialized Nsp-labeled ER domains, and both co-localized well with BFP-KDEL and with BFP-Sec61β throughout a “typical” ER network (Figure 1D and E, top and middle panels). In contrast, Nsp6 expression was sufficient to generate Nsp6-labeled domains. Notably, Nsp6 expression did not disrupt the overall structure of the general ER when visualized using the BFP-KDEL marker (Figure 1D, bottom left panel). However, the Nsp6 domains had two striking qualities that resembled results obtained when all three Nsps were expressed: they excluded BFP-KDEL and were highly enriched for Sec61β (compare Figure 1E, bottom right panel to Figure 1C).

Co-expression of β-coronavirus Nsp3 with Nsp4 has been shown to induce the formation of DMV-like ER membrane structures by EM in transfected cells^9–13,25^. Therefore, we asked whether co-expression of MHV Nsp3 and Nsp4 together in FL83B cells also generated specialized Nsp3/4-labeled ER domains. FL83B cells were co-transfected with mNe-Nsp3 and mCh-Nsp4 and either BFP-KDEL or BFP-Sec61β (Figure 1F and 1G, respectively). As predicted^9,11,25^, expression of Nsp3 and Nsp4 together generated domains that excluded BFP-KDEL, indicative of zippering (Figure 1F, in yellow boxes). However, these Nsp3/4 domains did not enrich or alter the ubiquitous localization of BFP-Sec61β (Figure 1G). In contrast, any double co-transfection that included Nsp6 and either Nsp3 or Nsp4 produced Nsp6-positive ER domains that were enriched with Sec61β (Supplementary Figure 1A and B). In these double transfections, we also observed re-localization of Nsp4, but not Nsp3, from the general ER to “wrap” Nsp6 domains, indicating Nsp3 re-localization to Nsp6 domains relied on Nsp4 expression, likely through Nsp3 and Nsp4 luminal interactions^25^ (Supplementary Figure 1A, compare mNe-Nsp3 to mCh-Nsp4, 1B). Notably, expression of mNe-Nsp3, mCh-Nsp4, and HALO-Nsp6 together in Cos-7 cells also produced Nsp3/4/6 domains that sequestered BFP-Sec61β and excluded BFP-KDEL (Supplementary Figure 1C and D). As before, expression of just Nsp3, Nsp4 or Nsp6 in Cos-7 cells did not disrupt the homogenous ER distribution of BFP-KDEL (Supplementary Figure S1E). Nsp6 expression alone was also sufficient in Cos-7 cells to generate Nsp6 domains that sequestered BFP-Sec61β (Supplementary Figure S1F). Together, our data suggest that Nsp6 is capable of both remodeling ER membranes into Nsp6 domains and sequestering Sec61β into these domains in multiple cell types.

Previous work has implicated ER-shaping proteins in β-coronavirus DMV biogenesis^26,27^. Nsp6 remodeled ER domains are unique in their morphology. We therefore wondered if they would either be enriched with ER proteins that localize exclusively to ER tubules or with ER proteins that are enriched in rough ER sheets^28,29^. To visualize Nsp6 domains relative to the tubular ER, FL83B cells were co-transfected with Halo-Nsp6 (to form Nsp6 domains, magenta) and the tubular ER protein mNe-Rtn3L (Reticulon 3L)^28^. Rtn3L was not recruited into Nsp6 domains (Figure 1H). Conversely, we asked whether Nsp6 domains are derived from ER sheets by co-transfecting FL83B cells with Halo-Nsp6, BFP-Sec61β and the sheet-shaping protein mNe-SigmaR1^29^. We again saw no observable recruitment of mNe-SigmaR1 into Nsp6 domains (Figure 1I). Finally, we visualized Nsp6 domains relative to Climp63, an ER protein that localizes to both peripheral ER tubules and sheets where it functions to regulate lumenal spacing^30,31^. Co-transfection of FL83B cells with Halo-Nsp6, BFP-KDEL, and mCh-Climp63 also showed that Climp63 was absent from Nsp6 domains (Figure 1J). Together, these data reveal that Nsp6 membrane structures do not adopt a conformation facilitating co-localization with three major ER-shaping proteins.

### Nsp6 domains selectively sequester the Sec61 translocon

We were intrigued by the ability of Nsp6 to sequester Sec61β out of the general ER and into Nsp6 domains and wanted to validate this observation quantitatively. We imaged FL83B cells transiently transfected with mNe-Sec61β (green) and mCh-KDEL (magenta), with or without HALO-Nsp6 (blue) and performed Manders’ colocalization coefficient (MCC)^32^ analyses on 10×10 µm regions of interest (ROIs) near the periphery of cells where the ER network was well resolved (Figure 2A, see Control). As a control, one channel was rotated 90° prior to MCC analysis to simulate a random overlap score (Control 90°). In control conditions, mNe-Sec61β co-localized well with mCh-KDEL (Figure 2B, Control MCC = 0.88) and this score was reduced following a 90° rotation (Figure 2B, Control 90° MCC = 0.43). In contrast, mNe-Sec61β did not co-localize with mCh-KDEL when Nsp6 is present (Figure 2B, MCC = 0.45) because it was concentrated into Nsp6 domains (Figure 2C, MCC = 0.89), which both qualitatively and quantitatively exclude the KDEL luminal protein (Figure 2D MCC = 0.36). Together, these analyses demonstrate that Sec61β was depleted from the general ER because it had been sequestered into Nsp6 domains.

**Figure 2:**
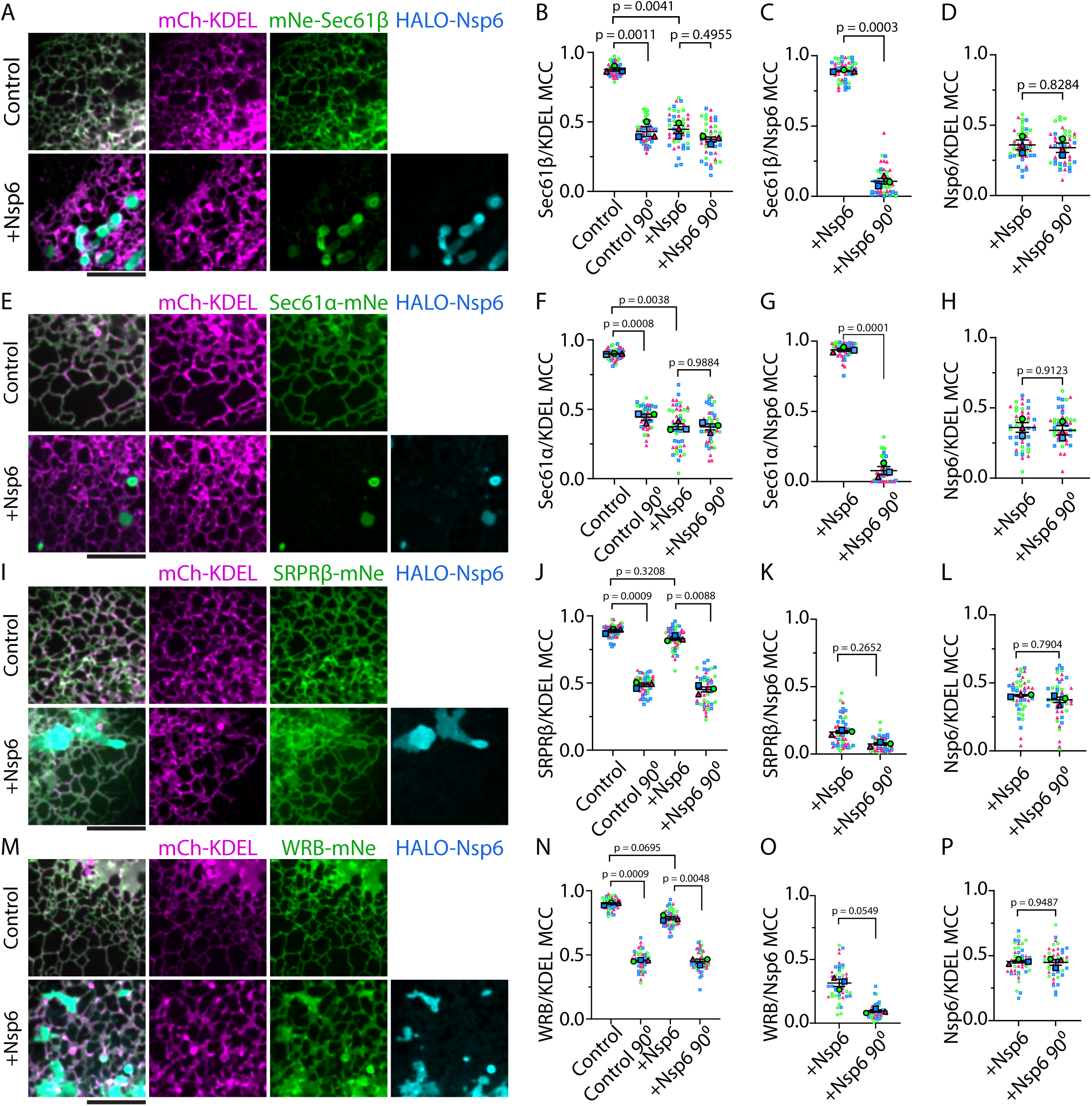
Nsp6 domains selectively sequester the Sec61 translocon. Representative images of FL83B cells co-transfected with (A) mCh-KDEL (magenta), mNe-Sec61β (green), and in the absence (top panels) or presence (bottom panels) of HALO-Nsp6 (JFX650 conjugated, blue). (B-D) A 10 x 10 µm ROIs were used to score protein co-localization (between channels indicated) within ER domains by Mander’s Correlation Coefficient (MCC) analyses. Control 90° and +Nsp6 90° are the MCC measured after rotating one channel 90° relative to the other channel to account for overlap due to chance and pixel density. (E) Images of FL83B cells co-transfected with mCh-KDEL (magenta), Sec61α-mNe, and in the absence (top panels) or presence (bottom panels) of HALO-Nsp6 (blue). (F-H) MCC were measured and graphed as in (B-D) for (E). (I) Images of FL83B cells co-transfected with mCh-KDEL (magenta), SRPRβ-mNe, and in the absence (top panels) or presence (bottom panels) of HALO-Nsp6 (blue). (J-L) MCC were performed as in (B-D) for (I). (M) Images of FL83B cells co-transfected with mCh-KDEL (magenta), WRB-mNe, and in the absence (top panels) or presence (bottom panels) of HALO-Nsp6 (blue). (N-P) MCC were measured and graphed as in (B-D) for (M). For MCC graphs, the large blue square, green circle, and magenta triangle represent the mean for independent biological replicates and corresponding small symbols are each individual data point within the respective replicate (3 biological replicates, n = 15 cells per replicate, error bars = ± standard error of the mean). Scale bars = 5 µm.

Sec61 is the core component of the ER translocation channel or “translocon” which is bound by ribosomes and functions to co-translationally translocate nascent polypeptide chains either across or into the ER membrane^14–18^. Sec61β is a nonessential subunit found in complex with the main pore-forming subunit, Sec61α^33^. The Sec61β subunit has been extensively used as a reliable membrane marker for scoring ER morphology^28,34–36^. We were surprised to see the degree to which this subunit was qualitatively hijacked from the general ER into Nsp6 domains. To determine if this was true for other Sec61 components, we next imaged Sec61α-mNeonGreen (green) an essential subunit that forms the central channel of the translocon that is required for ribosome docking and protein translocation^17,18,37^. FL83B cells were transiently co-transfected with Sec61α-mNe (green) and mCh-KDEL (magenta) and with or without HALO-Nsp6 (blue). In the absence of Nsp6, Sec61α-mNe co-localized well with mCh-KDEL throughout the ER network both qualitatively and quantitatively (Figure 2E and F, Control MCC = 0.90; reduced to Control MCC 90° = 0.44 after a 90° rotation). However, in the presence of Nsp6, Sec61α was sequestered into Nsp6 domains (Figure 2G; MCC = 0.94 vs 0.08 after 90° rotation) and no longer co-localized with mCh-KDEL (Figure 2H; MCC = 0.37 vs 0.37 after 90° rotation). Thus, the sequestration of both Sec61α and Sec61β into Nsp6 domains is accompanied by their corresponding depletion from the general ER. Notably, we took care to express exogenous Sec61a-mNe protein (upper arrowhead) at levels that are comparable to endogenous Sec61α (lower arrowhead) according to western blot analysis (Supplementary Figure 2A).

The signal recognition particle (SRP) and its receptor function to target ribosomes and their associated transcripts to the Sec61 translocon for co-translational translocation^38–40^. We next assessed whether the receptor, SRPRβ, which binds to and functions upstream of the Sec61 translocon, was also sequestered into Nsp6 domains. We visualized the localization of the membrane bound subunit of the SRP receptor (SRPRβ-mNe, green) relative to mCh-KDEL (magenta) in the absence or presence of HALO-Nsp6 (blue) in FL83B cells. SRPRβ-mNe co-localized qualitatively and quantitatively with mCh-KDEL throughout the general ER and this did not change upon co-expression of Nsp6 (Figure 2I and J, MCC = 0.89 vs 0.83, respectively). In contrast to Sec61, SRPRβ-mNe was neither sequestered into nor excluded from Nsp6 domains (Figure 2K-L).

The Guided entry of tail-anchored protein (GET) complex is an alternative targeting pathway that functions post-translationally to insert tail-anchored proteins into the ER^41,42^. We imaged a membrane bound component of the GET complex WRB-mNe (aka GET1) and mCh-KDEL with and without co-transfection of Nsp6 in FL83B cells to visualize if the GET complex would be sequestered into Nsp6 domains. Like SRPRβ, WRB-mNe co-localized well with the mCh-KDEL marker throughout the general ER both qualitatively (Figure 2M) and quantitatively (Figure 2N control MCC = 0.90 vs 0.79, respectively) regardless of whether Nsp6 domains were present. And like SRPRβ, WRB-mNe was neither sequestered by nor excluded from Nsp6 domains (Figure 2M-P). Together, these data reveal that co-translational and post-translational targeting machineries retain access to Nsp6 domains but are not specifically retained there.

### Nsp6 domains sequester the multipass membrane protein insertases

We sought to identify whether Nsp6-remodeled ER domains sequester other ER proteins that function to co-translationally translocate membrane proteins. We performed proximity biotinylation experiments by fusing the promiscuous biotin ligase TurboID^43^ to mNe-Nsp6. FL83B cells were transiently transfected with TurboID-mNe-Nsp6, a TurboID-mNe cytosolic control, or a mock transfection control, and incubated with 500 µM of biotin for 1 hour. Immuno-staining with AlexaFluor 405-conjugated streptavidin antibody (green) confirmed that Nsp6 domains were enriched with biotinylation signal (Figure 3A, see +Nsp6 in magenta) compared to the cytosolic control, which showed diffuse staining (Figure 3A). Cell lysates were collected and biotinylated proteins were purified on a streptavidin affinity column and identified by mass-spectrometry (Figure 3B). The iBAQ values for the TurboID-mNe-Nsp6 samples were divided by the iBAQ values for the TurboID-mNe cytosolic or mock transfection control (whichever iBAQ value was highest) to increase the likelihood of identifying relevant ER proteins that are biotinylated within Nsp6 domains (see Methods). Nsp6 self-biotinylation was the top hit (Figure 3C, blue dot). The top 30 ER protein candidates biotinylated by TurboID-mNe-Nsp6 are listed in order of enrichment (black dots, also see Supplementary Table 1). Biotinylated ER protein hits that are subunits of translocation machineries (i.e. EMC4 and OPTI) are indicated with protein names (Fig 3C, magenta dots). ER hits that are known to be translocation-associated factors involved in post-processing of inserted proteins are also highlighted (Fig 3C, green dots).

**Figure 3:**
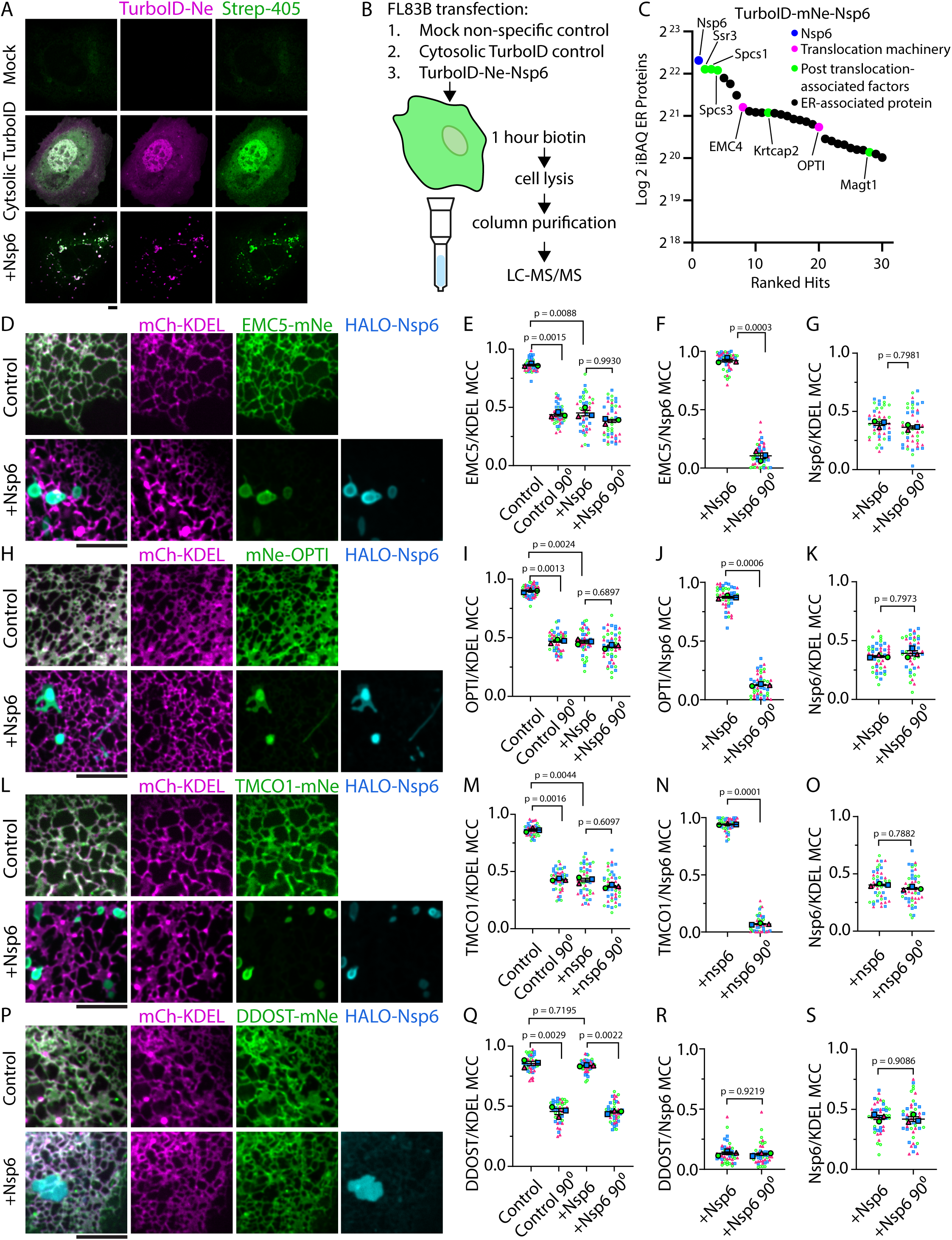
Nsp6 domains sequester the multipass membrane protein insertases. (A) Representative images of FL83B cells (top) mock transfected, (middle) transfected with TurboID-mNe empty vector or (bottom) TurboID-mNe-Nsp6 (magenta) and treated with biotin for 1 hour prior to immunostaining with streptavidin-405 antibody (green). (B) TurboID biotinylation experimental approach. (C) Log_2_ iBAQ values plotted for the top 30 ER-specific biotinylation hits identified by mass-spectrometry. Nsp6 self-biotinylation (blue dot), translocation machineries (magent dots), post-translocon-associated factors (green dots), and all other ER proteins (black dots). Representative 10 x 10 µm images of (D) EMC5-mNe (H) mNe-OPTI, (L) TMCO1-mNe, or (P) DDOST-mNe (green) and mCh-KDEL (magenta) co-transfected in the absence (top) or presence of HALO-Nsp6 (bottom, blue). MCC analysis of EMC5-mNe, (I) mNe-OPTI, (M) TMCO1-mNe, or (Q) DDOST-mNe with mCh-KDEL without (Control) and with (+Nsp6) HALO-Nsp6 including an mCh-KDEL channel rotated control (Control 90° and +Nsp6 90°). MCC analysis of (E) EMC5-mNe, (I) mNe-OPTI, (M) TMCO1-mNe, or (Q) DDOST-mNe with HALO-Nsp6 (+Nsp6) compared to HALO-Nsp6 channel rotated control (+Nsp6 90°). MCC analysis of (G, K, O, S) HALO-Nsp6 with mCh-KDEL (+Nsp6) and mCh-KDEL rotated control (+Nsp6 90°). For MCC graphs, the large blue square, green circle, and magenta triangle represent the mean for independent biological replicates and corresponding small symbols are each individual data point within the respective replicate (3 biological replicates, n = 15 cells per replicate, error bars = ± standard error of the mean). Scale bars = 5 µm.

A subunit of the ER Membrane Complex (EMC), EMC4, was biotinylated by TurboID-mNe-Nsp6. The EMC is composed of 9 subunits (7 of which are membrane spanning) and functions in concert with Sec61 as a chaperone for multipass TMD topogenesis and as a solo insertase^44,45^. First, we visualized mNe-EMC4 (blue) relative to mCh-KDEL (green) in the presence of HALO-Nsp6 domains (magenta) in FL83B cells. EMC4 was qualitatively enriched in Nsp6 domains (Supplementary Figure 2B). Next, we tested if the EMC would be sequestered by Nsp6 domains by co-transfecting FL83B cells with mCh-KDEL (magenta) and with a membrane-spanning subunit of the EMC that is required for EMC assembly and stability (EMC5-mNe, in green)^46^ with or without HALO-Nsp6 (blue). Live images showed EMC5 co-localized well with mCh-KDEL in the absence of Nsp6 (Figure 3D and E control MCC = 0.86). However, like the Sec61 translocon, the EMC5-mNe insertase was qualitatively and quantitatively sequestered out of the general ER and into HALO-Nsp6 domains (Figure 3F MCC = 0.92), which exclude mCh-KDEL (Figure 3G and E). Western blot analysis confirmed that the ectopically expressed EMC5-mNe (upper arrowhead) was expressed at lower levels than endogenous EMC5 (lower arrowhead, Supplementary Figure 2C).

The GET and EMC-like (GEL) complex is an ER insertase for the MPT (multi-pass membrane protein translocon) that interacts with the Sec61 translocon^47–49^. The GEL complex is composed of subunits OPTI and TMCO1^45^. OPTI was biotinylated by TurboID-mNe-Nsp6 (Figure 3C). To test whether the GEL complex would be sequestered into Nsp6 domains, we transiently co-transfected FL83B cells with mCh-KDEL and either mNe-OPTI or TMCO1-mNe in the absence or presence of HALO-Nsp6. Both GEL subunits co-localized well with the general ER mCh-KDEL marker in the absence of Nsp6 (Figure 3H, I, and L, M MCC = 0.90 and 0.86). Like the Sec61 translocon and the EMC insertase, the GEL complex components were also qualitatively and quantitatively sequestered into HALO-Nsp6 domains (Figure 3J and 3N OPTI/Nsp6 and TMCO1/Nsp6 MCC = 0.97 and 0.94, respectively) and were depleted from the general ER (Figure 3I and M +Nsp6 MCC = 0.47 and 0.43, respectively).

TurboID-mNe-Nsp6 also biotinylated accessory proteins that are associated with the Sec61 translocon but do not function directly in translocation. Two noncatalytic subunits of the Oligosaccharyltransferase (OST) complex, KRTCAP2 (OST-A complex) and MAGT1, Magnesium Transporter 1 (OST-B complex)^50–52^ were biotinylated by TurboID-mNe-Nsp6. The OST complex interacts with the Sec61 translocon to N-link or O-link glycosylate nascent chains as they are co-translationally translocated (N-linked, OST-A) or post-translationally (N- or O-linked, OST-B)^53,54^. To visualize whether the OST complex would be recruited to Nsp6 domains, FL83B cells were co-transfected with mCh-KDEL and an OST complex component DDOST-mNe (Disorders of the OST), an essential core subunit of both OST-A and OST-B^55^, with and without HALO-Nsp6. DDOST co-localized well with mCh-KDEL regardless of whether Nsp6 domains were present (Figure 3P and Q, Control MCC = 0.86 vs MCC = 0.84, respectively). Like mCh-KDEL, DDOST-mNe was quantitatively excluded from Nsp6 domains (Fig 3R and S, DDOST/Nsp6 MCC = 0.13). It is not surprising that the OST complex would be excluded from Nsp6 domains because the GEL and OST complex occupy overlapping and thus mutually exclusive positions behind the Sec61 lateral gate^56,57^. Furthermore, co- and post-translational N- and O-glycosylation required for the maturation of β-coronaviruses viral structural proteins S, E, and M, occurs at the ER-Golgi intermediate compartment (Ergic)^58–64^. Indeed, we can visualize that Nsp6 domains are nearby but not overlapping with Ergic components in our transfected FL83B cells (Supplementary Figure 2D).

Other translocon-associated accessory proteins that were biotinylated by TurboID-mNe-Nsp6 include Ssr3 (TRAP-γ) a subunit of the Signal Sequence Receptor complex (TRAP complex) and Spcs1 (Spc12) and Spcs3 (Spc22/23); two subunits of the Signal peptidase complex (SPC, Figure 3C, green dots). The TRAP is associated with the Sec61 translocon via Ssr3 (TRAP-γ)^65^ and can facilitate the insertion of certain membrane protein substrates^66,67^. Therefore, we visualized two subunits of the TRAP complex, mNe-Ssr3 and Ssr1-mNe (TRAP-α, blue), relative to mCh-KDEL (green) in the presence of HALO-Nsp6 domains (magenta) in FL83B cells. Ssr3 was qualitatively enriched at Nsp6 domains, whereas Ssr1 was not (Supplementary Figure 2E and F). These data indicate that only Ssr3, the TRAP complex subunit that is known to interact directly with the Sec61 translocon^65^ is likely sequestered into Nsp6 domains.

The SPC can interact transiently with the Sec61 translocon to cleave N-terminal signal sequences from nascent proteins as they are translocated by the Sec61 translocon^68,69^. We visualized three subunits of the SPC (Spcs1-mNe, Spcs2/Spc25-mNe, and mNe-Spcs3, blue) relative to mCh-KDEL (green) in the presence of HALO-Nsp6 domains (magenta) in FL83B cells. Only Spcs2 appeared enriched at Nsp6 domains, whereas Spcs1 and Spcs3 were not (Supplementary Figure 2G, H,and I). Interestingly, Spcs2 is also the component of the SPC complex that interacts directly with the Sec61 translocon^68,70^. Together, these data suggest that Nsp6 domains can recruit other ER proteins (from TRAP and SPC complexes) that interact with the Sec61 translocon, though not necessarily in complexed form. It is likely that some of the biotinylated hits we observed occurred during the initial translation and translocation of TurboID-mNe-Nsp6 but are not retained in Nsp6 domains following their formation.

### Nsp6 domains restrict translocon mobility

Our data revealed that Nsp6 expression remodeled the ER membrane into Nsp6 domains that sequestered the Sec61 translocon, the EMC, and the GEL multipass translocation machineries away from the general ER. In contrast, upstream components involved in co-translational or post-translation targeting to the ER were neither sequestered nor excluded from these Nsp6 domains. We performed fluorescence recovery after photobleaching (FRAP) experiments to test whether Nsp6 domains are indeed continuous with the ER and allow the free lateral diffusion of SRPRβ-mNe and Sec61α-mNe (in grey) into Nsp6 domains (magenta). First, we established a baseline for the rate of fluorescence recovery of SRPRβ-mNe and Sec61α-mNe expressed in cells that do not contain Nsp6 domains. Similar sized ROIs (white circles) were selected in the peripheral ER, photobleached, and recovery was recorded over time (Figure 4A and C). FRAP experiments reveal that SRPRβ-mNe and Sec61α-mNe had similar mobile fractions (70% vs. 72%) and similarly rapid rates of recovery back into photobleached ROIs (Figure 4A and C: t_1/2_ = 5.74 vs 8.59 sec and k = 0.12 vs 0.08 sec^-1^). Next, we performed FRAP experiments on SRPRβ-mNe or Sec61α-mNe in similar sized ROIs that overlapped with HALO-Nsp6 domains. The mobile fraction was much higher for SRPRβ-mNe (Figure 4B, 68%) than for Sec61α-mNe (Figure 4D, 18%), which indicated there was a smaller freely diffusive pool of Sec61α-mNe consequently limiting recovery in Nsp6 domains. However, the rates of fluorescence recovery into ROIs that overlap with Nsp6 domains were similar for SRPRβ-mNe (Figure 4B, graph t_1/2_ = 10.29 sec and k = 0.07 sec^-1^) and Sec61α-mNe (Figure 4D, graph t_1/2_ = 8.72 sec and k = 0.08 sec^-1^). These data confirmed that Nsp6 compartments were continuous with the ER and did not restrict SRPRβ and Sec61α from diffusing into Nsp6 domains.

**Figure 4:**
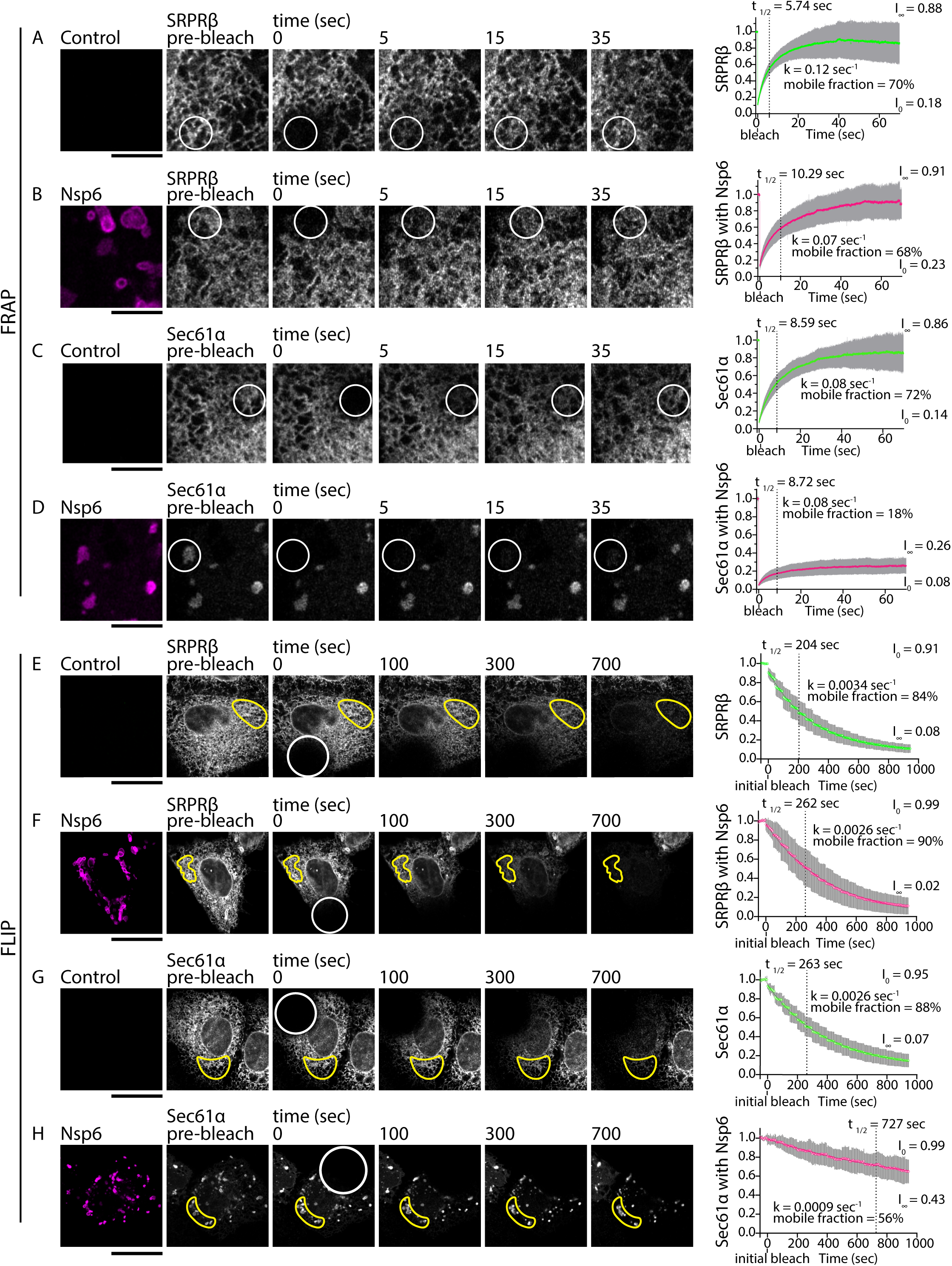
Nsp6 domains restrict translocon mobility. Representative time-lapse images of FL83B cells transfected with (A) SRPRβ-mNe (white) or (A) SRPRβ-mNe (white) and HALO-Nsp6 (magenta) imaged before (prebleach) and after photobleaching a region of interest (ROI) in circle shown. Fluorescence recovery was measured within the ROI at times indicated following the photobleaching event and were plotted (on the right as indicated). Statistics measured during fluorescence recovery after photobleaching (FRAP) as indicated. (C-D) Experiments were performed and recorded as in (A-B) for FL83B cells transfected with (C) Sec61α-mNe (white) or (D) Sec61α-mNe (white) and HALO-Nsp6 within the general ER network (Control) or within an Nsp6 domain (Nsp6, magenta), respectively. (E-F) FL83B cells were transfected with (E) SRPRβ-mNe (white) or (F) SRPRβ-mNe (white) and HALO-Nsp6 (magenta) and were bleached in the white circle ROIs beginning at time = 0 and again every 50 seconds. Fluorescence loss in photobleaching (FLIP) was measured in another region of the cell at ROIs (yellow outline) that corresponded to control or Nsp6 domains. FRAP and FLIP curves were fit using a one phase association (FRAP) or one phase decay (FLIP) to extrapolate the mobile fraction, half-time recovery or loss (dashed line, t_1/2_), recovery or decay rate (k), normalized intensity at t = 0 (I_0_), and plateau (I_∞_). Means for each time-point are shown with green (control) or magenta (+Nsp6) circles (3 replicates, n = 5 cells per replicate, error bars = ± standard deviation). Scale bars = 5 µm (A-D). or 20 µm (E-H).

Next, we performed complementary fluorescence loss in photobleaching (FLIP) experiments to compare the ability of SRPRβ and Sec61α to diffuse out of Nsp6 domains. FLIP experiments were performed on FL83B cells expressing SRPRβ-mNe or Sec61α-mNe with and without HALO-Nsp6. A large ROI placed in the peripheral ER was re-bleached every 50 seconds (white circle) and the rate of fluorescence loss in an unbleached ROI (yellow) was measured. The mobile fraction for SRPRβ-mNe and Sec61α-mNe were similar in control cells (Figure 4E and G mobile fractions = 84% and 88%). The rate of fluorescence loss within the unbleached ROI was also similar between SRPRβ-mNe and Sec61α-mNe in control cells (Figure 4E and G graphs k = 0.0034 vs 0.0026 sec^-1^ and t_1/2_ = 204 sec vs. 263 sec). In cells with HALO-Nsp6, the statistics are similar for SRPRβ-mNe in an ROI that overlaps HALO-Nsp6 domains (yellow) compared to SRPRβ-mNe in control cells (Figure 4F, mobile fraction = 90%, k = 0.0026 sec^-1^ and t_1/2_ = 262 sec), indicating that lateral diffusion out of Nsp6 domains was not restricted. In stark contrast, Sec61α-mNe fluorescence within an Nsp6 domain (yellow) was qualitatively and quantitatively observed throughout the FLIP experiment and exhibited dramatically slowed fluorescence loss with continuous bleaching (Figure 4H, note Sec61α signal at 700 seconds, mobile fraction = 56%, k = 0.0009 sec^-1^ and t_1/2_ = 727 sec). Taken together, our FRAP and FLIP experiments demonstrate that Nsp6 domains are continuous with the ER but selectively retain Sec61α-mNe.

### MHV replication centers recruit Sec61β

We next aimed to test whether FL83B cells infected with MHV similarly sequester Sec61β near DMVs. First, we tested the ability of FL83B cells to support MHV strain JHM^IA^ viral infection. FL83B cells were infected with MHV JHM^IA^ (MOI of 0.1), cell pellets were collected, and RNA was isolated at specific timepoints following infection (see Figure 5A). We performed RT-qPCR to measure Nsp6 transcript levels in infected cell pellets (see Figure 5A step (B)). We saw a substantial change in the mean ΔCt values for infected cell pellets from 13.5 to 2.3 at 24 hpi which corresponded to a 3,401-fold increase in intracellular viral RNA compared to the 0 hpi control (Figure 5B, green), indicating MHV JHM^IA^ successfully replicated in FL83B cells within our time course. To compare viral Nsp6 transcript levels with HALO-Nsp6, FL83Bs were transfected with 500 ng of HALO-Nsp6 (as in Figures 1-2) or with a mock transfection control (black dot) which were processed along with the infected cell pellets for qPCR. The mean ΔCt value for transfected Nsp6 was 5.9 (magenta), which lay between the ΔCt mean values of cell culture pellets at 8 and 12 hpi (Figure 5B; mean = 8.5 and 3.9, respectively). These data demonstrate that Nsp6 transcript levels produced following transfection of the Nsp6 transgene are comparable to those observed during viral replication.

**Figure 5:**
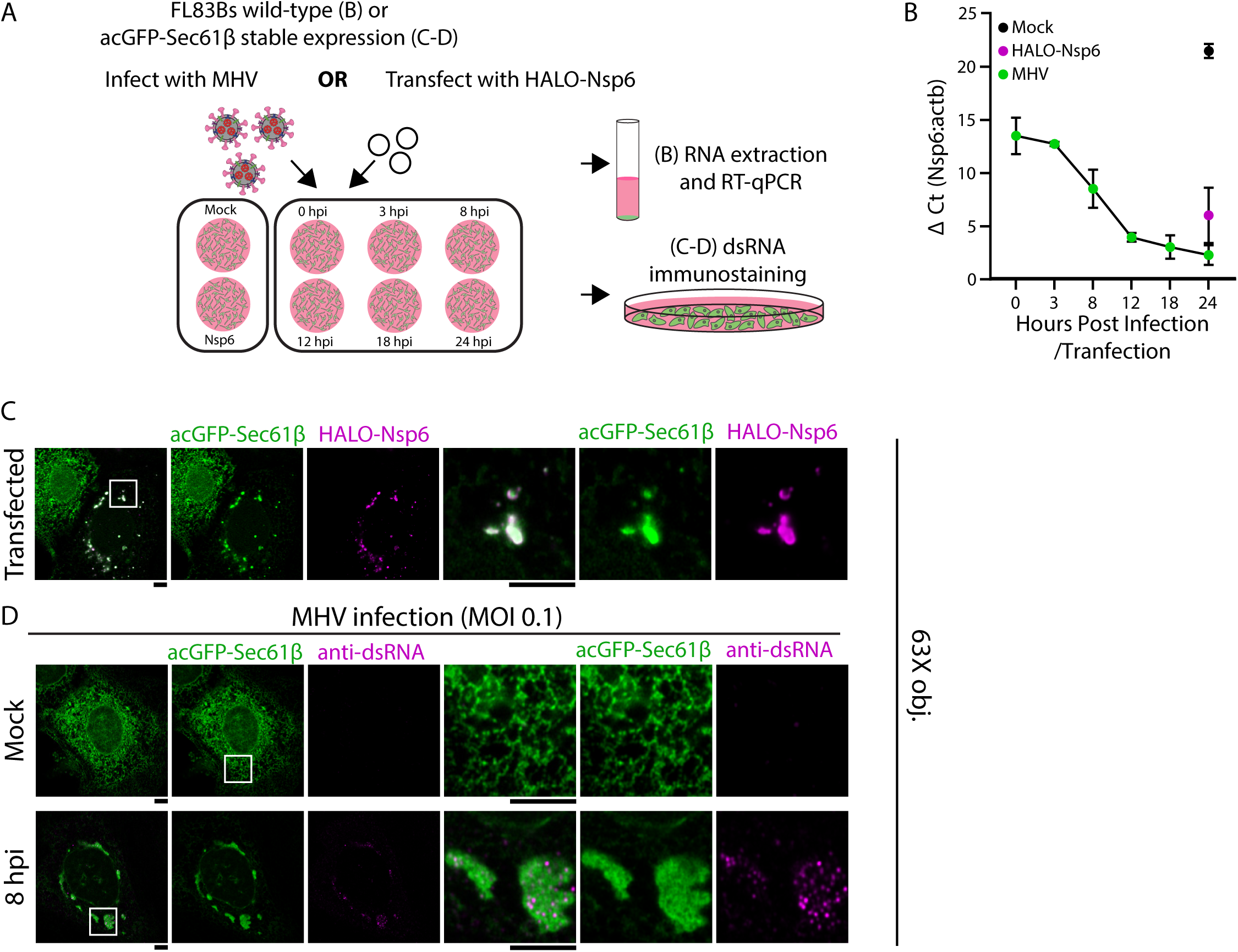
MHV replication centers recruit Sec61β. (A) Cartoon of experimental workflow performed for B-D. (B) FL83B cells or (C and D) FL83B cells stably expressing acGFP-Sec61β were transfected with (B and C) HALO-Nsp6 or (B and D, E) infected with 0.1 MOI MHV JHM^IA^ and collected at time points indicated for (B) qPCR or (C and D) immunostaining analysis. (B) qPCR plot of ΔCt (Ct, cycle threshold) values calculated from the difference between Nsp6 and actin (actb – 5, see Methods) in samples extracted from infected cell culture pellets (green), HALO-Nsp6 transfected (magenta), or mock transfected cells (black). (C and D) Representative 63X objective images of fixed FL83B cells stably expressing acGFP-Sec61β (green) and (C) HALO-Nsp6 transfected (JFX650 conjugated, 500 ng, magenta) or (D) 0.1 MOI MHV JHM^IA^ infected (mock control, 8, 12, and 18 hpi). Infected cells were immuno-stained with dsRNA to identify replication centers (magenta) relative to Sec61β signal (green). Scale bars = 5 µm (C, D).

Next, we generated FL83B cells stably expressing acGFP-Sec61β to test whether MHV infection would similarly alter Sec61β localization. Transfection of HALO-Nsp6 (magenta) into the acGFP-Sec61β stable cell line caused acGFP-Sec61β (green) to be sequestered into HALO-Nsp6 domains as previously scored (Figure 5C). Syncytial formation seen at 12 hpi and later time points made the fluorescence images difficult to assess, so we imaged DMVs formed in cells stably expressing acGFP-Sec61β at the 8 hpi timepoint prior to syncytia formation (Figure 5D). Infection led to a dramatic re-localization of acGFP-Sec61β (green) to discrete domains that resembled DMVs (Figure 5D)^71^. Cells were simultaneously immuno-stained with an antibody against double-stranded RNA (dsRNA, clone rJ2, magenta) which uniquely labels active viral replication sites^71^. DsRNA/DMV positive foci localization coincided with membrane domains containing sequestered acGFP-Sec61β (Figure 5D, 8 hpi 64X and 20X magnification in Supplementary Figure 3A). Together these data reveal that MHV infection sequesters Sec61β into active DMVs, recapitulating the pattern we observed in Nsp6-transfected FL83B cells.

Positive-strand RNA viruses are known to remodel host endomembranes to create replication centers that recruit specific host proteins required for viral replication^10,26,27,72–79^. Here, we show that membrane remodeling by the MHV non-structural protein Nsp6 generates membrane-bound compartments that sequester ER insertases, including the Sec61 translocon and two associated complexes that are involved in multi pass membrane protein insertion (EMC and GEL). The selective retention of insertases, coupled with the free diffusion of upstream targeting and accessory factors, indicates that Nsp6 does not restrict access of most general ER-resident membrane proteins, but does create a localized environment rich in insertases. It is still unclear how Nsp6 drives sequestration of translocons, either through direct binding with host translational factors or indirectly by morphologically re-shaping ER membrane structure^10^, These membrane domains—which are unique from ER-tubules or sheets—may function to promote local insertion or translocation of viral membrane proteins through hijacking of host insertases into discrete domains. Indeed, Nsp6 has been shown to cluster DMVs generated by Nsp3 and Nsp4^10^. Notably, we saw a clear rearrangement of Nsp3 and Nsp4 to “wrap” Nsp6 domains in an Nsp4-dependent manner, consistent with the notion that insertase-rich Nsp6 domains could promote local insertion of Nsp4, with Nsp3 recruitment to sites for DMV formation through interaction with Nsp4.

These observations support a model in which Nsp6 performs a dual role in replication organelle biogenesis—first reshaping membranes to generate a suitable scaffold and then recruiting host machinery required for viral membrane protein insertion. A similar reliance on ER insertases has been reported for other positive-strand RNA viruses. For example, Dengue virus NS4B is a multipass non-structural protein that, together with NS4A, drives ER remodeling to form viral replication compartments (vRCs), a process dependent on EMC-mediated insertion of both proteins. NS4B interacts with EMC4, which co-localizes with vRCs, linking EMC recruitment to flavivirus vRC formation^77,78^. By analogy, the enrichment of EMC, MPT, and Sec61 insertase subunits within Nsp6 domains might facilitate the insertion of viral membrane proteins and/or be a viral strategy to sequester this machinery away from host substrates. Although the mechanism by which Nsp6-mediated redistribution of insertases remains unresolved, these parallels suggest that coronaviruses, may employ a strategy analogous to flaviviruses, in which the host insertase machinery is repurposed to ensure efficient insertion or translocation of viral and host membrane proteins within virally induced ER domains.

## ACKNOWLEDGEMENTS

We thank AJ Sariol, SA Lowery, and NA Schuster, for generously sharing pBAC-JHM^IA^, virus stocks, and protocols for propagating virus and providing valuable technical advice. We thank TS Nahreini in the Flow Cytometry Core Facility for sorting the FL83B stable cell lines and S Maurya for technical advice on the mass-spectrometry results. We thank S Anderson, T Rizo-Garza, and S Zaganelli for help and discussions. SP was supported by grant R01 AI129269. GKV is an investigator of the Howard Hughes Medical Institute.

## METHODS

### Plasmids

pBAC-JHM.IA^80^ was used to PCR amplify (Invitrogen Platinum SuperFi II DNA Polymerase #12369010) non-structural protein genes Nsp3, Nsp4, and Nsp6 for Gibson Assembly (NEB #E2611L) into mammalian cell transient expression vectors. All constructs generated for this study include a short Gly-Gly-Gly-Ser linker between the fluorescent tag and target protein. Nsp3 was assembled into pmNeonGreen-CT vector (Allele Biotechnology) to generate a N-terminal mNeonGreen tagged Nsp3 construct (mNe-Nsp3). The mCherry gene was PCR amplified from pmCherry-C1 vector (Clonetech) and assembled into the backbone of pmNeonGreen-CT vector (excluding the mNeonGreen) with Nsp4 to generate a N-terminal mCherry tagged Nsp4 construct (mCh-Nsp4) with the same expression backbone as mNe-Nsp3. The HALO gene was PCR amplified from HALO-C1 vector and cloned into the backbone of pmNeonGreen-CT vector (excluding the mNeonGreen) with Nsp6 to generate a N-terminal HALO tagged Nsp6 construct (HALO-Nsp6) with the same expression backbone as mNe-Nsp3. BFP-Sec61β (Addgene #49154)^36^, BFP-KDEL (Addgene #49150), and mCherry-KDEL have been previously described^81^. cDNA from mouse liver tissue (Biochain #C1334149) was used as a template to PCR amplify Sec61α, Sec61β, SRPRβ, WRB (GET1), EMC4, EMC5, DDOST, TMCO1, OPTI, Spcs1 (Spc12), Spcs2 (Spc25), Spcs3 (Spc22/23), Ssr3 (TRAP-γ), and SSR1 (TRAP-α) which were also Gibson assembled into pmNeonGreen-CT or - NT vectors (Allele Biotechnology) to generate Sec61α-mNe, mNe-Sec61β, SRPRβ-mNe. WRB-mNe, mNe-EMC4, EMC5-mNe, DDOST-mNe, mNe-OPTI, TMCO1-mNe, Spcs1-mNe, Spcs2-mNe, mNe-Spcs3, mNe-Ssr3, and SSR1-mNe. mCh-Climp63 (Addgene #136293)^82^, mNe-SigmaR1 (Addgene #226566)^29^, and mNe-Rtn3L (Addgene #169034)^83^ in Figure 1 and GFP-Ergic53 (a gift from the Hans Peter-Hauri lab) in Supplementary Figure 2 have been previously described. TurboID-mNe-Nsp6 used in proximity labeling biotin ligase experiments were assembled by Gibson assembly with PCR amplified inserts TurboID-mNeonGreen (from pLV-TurboID-mNe)^43,84^ and Nsp6 (from HALO-Nsp6) into the PCR amplified vector backbone of pmNeonGreen-CT (excluding the inherent mNeonGreen). For the empty vector control TurboID-mNe, the Gibson assembly was performed with a PCR amplified insert of TurboID-mNeonGreen. The TurboID and mNeonGreen are separated by sequence Ala-Ala-Ala-Leu-Glu-Leu-Val-Asp-Pro linker. For generating FL83B stable cell lines in Figure 5, pLV-acGFP-Sec61β lentiviral vector was used and has been previously described^84^. 3^rd^ generation lentiviral helper plasmids are previously described and are available on Addgene: pCMV-VSV-G (Addgene #8454), pRSV-REV (Addgene #12253), and pMDLg-RRE (Addgene 12251).

### Cell lines

FL83B cells were purchased from ATCC (CRL-2390) and are used in all figures of this study apart from Supplementary Figure 1, which was done in COS-7 cells (ATCC CRL-1651). 293T cells (ATCC CRL-3216) were used to produce lentiviral particles for generating acGFP-Sec61β FL83B stable cells. 17Cl-1 and HeLa-MHVR^85^ cells were from Dr. Stanley Perlman and were used to passage MHV (17Cl-1) and titer the virus (HeLa-MHVR).

### Cell culture and transfection

All cell lines were maintained at 37⁰C and 5% CO_2_. FL83B and acGFP-Sec61β-FL83B cells were grown in Ham’s F-12K (Kaighn’s) Medium (F-12K/Gibco 21127022) and COS-7, HeLa-MHVR, and 293T cells were grown in Dulbecco’s Modified Eagle Medium (DMEM/Gibco 12430062), each supplemented with 10% heat-inactivated fetal bovine serum (Sigma F2442) and 1% penicillin-streptomycin (Gibco 15140122). 293T cell growth media was additionally supplemented with 1x GlutaMAX (Gibco 35050079). 17Cl-1 cells were grown in DMEM (Gibco 12430062) supplemented with 5% fetal bovine serum, 5% tryptose phosphate broth (Thermo 18050039), and 1% penicillin-streptomycin.

For fluorescence microscopy experiments with FL83Bs and COS-7, 220,000 FL83B cells or 200,000 COS-7 cells (counted with Bright-Line™ Hemacytometer/Sigma Z359629) were seeded on a 35-mm glass bottom dish (Cellvis D35-20-1.5-N) in 2 mL of growth media the day before to achieve ∼70% confluence on the day of transfection with plasmid DNA. FL83B cells were transfected with an optimal ratio of 1:4 plasmid DNA to FuGENE® 4K Transfection Reagent (Promega E5911). Lipid-DNA complex formation was allowed to occur for 20 minutes in Opti-MEM™ I Reduced Serum Medium (Gibco 31985070) prior to direct addition to cell growth media following the manufacturer’s protocol. For immunoblotting experiments, 1.5 x 10^6^ FL83B cells were seeded per 10-cm tissue culture dish (CELLSTAR® 82050-916) to achieve ∼70% confluency on the day of transfection. Transfection volumes and DNA plasmid amounts were scaled up according to surface area (6.8x 35-mm dish) following manufacturer’s instructions. For lentiviral particle production, 1.2 x 10^6^ 293T cells were seeded per well in a 6-cm dish to achieve ∼80% confluency on the day of transfection. COS-7 and 293T cells were transfected with Lipofectamine 3000 (Thermo Fisher L3000015) following the manufacturer’s protocol using an optimal plasmid DNA:P3000 ratio of 1:2 and complex incubation time of 20 minutes prior to direct addition to cell growth media.

The following concentrations of plasmid DNA were transfected in FL83B cells seeded in 35-mm dishes or 6-well plate: 2000 ng mNe-Nsp3; 1000 ng Sec61α-mNe, SRPRβ-mNe, mNe-WRB-mNe, DDOST-mNe, TMCO1-mNe, Spcs1-mNe, Spcs2-mNe, Spcs3-mNe, SSR1-mNe; 800 ng mCh-Nsp4, BFP- and mNe-Sec61β, mNe-EMC4, EMC5-mNe, mNe-OPTI, mCh-Climp63, mNe-SigmaR1, mNe-Rtn3L, GFP-Ergic53; 500 ng TurboID-mNe and HALO-Nsp6, TurboID-mNe-empty vector, BFP- and mCh-KDEL. FL83B cells seeded in 10-cm dishes: 6,800 ng Sec61α-mNe and TMCO1-mNe; 5,440 ng EMC5-mNe; 3,400 ng TurboID-mNe- and HALO-Nsp6, and TurboID-mNe-empty vector. COS-7 cells: 700 ng mNe-Nsp3; 300 ng mCh-Nsp4 and BFP-Sec61β; 200 ng HALO-Nsp6 and BFP-KDEL. 293T cells for lentiviral particle production: 4.5 µg pLV-acGFP-Sec61β, 4.0 µg pMDLg-RRE, 2.0 µg pRSV-REV and pCMV-VSV-G.

### Mouse hepatitis virus passaging and plaque assay

Protocols for viral passaging in 17Cl-1 cells and the HeLa-MHVR plaque assay were provided by the Perlman laboratory^23,85,86^. For viral passaging of P1 stock virus, 17Cl-1s were seeded in T75 flasks to reach ∼80% confluence on the day of infection. A JHM^IA^ P1 stock was rapidly thawed and diluted in 3 mL of pre-warmed DMEM to infect a single T75 flask at an MOI 0.01. Cells were incubated in infectious media at 37⁰C for 30 minutes, with gentle rocking every 5 minutes. Following incubation, 8 mL of fresh, pre-warmed 17Cl-1 media was added to the cell flask and incubated for 20 hours. Following 20 hours, flasks were frozen at −80⁰C, thawed, and cell debris were clarified at 3000 x g for 10 minutes. Supernatants (P2 stocks) were aliquoted, flash frozen in liquid nitrogen, and stored at −80⁰C before use.

For plaque assays, HeLa-MVHR cells were seeded into two CELLSTAR® 12-well cell culture plates (Avantor 82050-928) the day before viral titering to achieve ∼80% confluency. A JHM^IA^ P2 stock was thawed and serially diluted in 1 mL DMEM (10^−1^, 10^−2^, 10^−3^, 10^−4^, 10^−5^, 10^−6^, 10^−7^). HeLa-MHVR media was aspirated, and 200 µL of each dilution, in duplicate, was added to each well (including a DMEM-only control). Plates were incubated at 37⁰C for 30 minutes with gentle rocking every 5-10 minutes. For agar plugs, sterile 1.2% agar (BD Bacto™ Dehydrated Agar/BD 214030) in 1x DPBS was melted in the microwave and mixed 1:1 (10 mL:10mL) with pre-warmed sterile plug media (1 pack DMEM powder/Gibco 12100046, 5% FBS, 16% sodium bicarbonate, 2x GlutaMAX, 2x MEM non-essential amino acid solution (Gibco 11140050), 2% penicillin-streptomycin, 2 mM sodium pyruvate (Thermo Fisher 11360070), 500 mL ddH2O) in a 50 mL conical and mixed by inversion. Prior to mixing, agar solution was left to cool in a 55⁰C water bath for 20 minutes. Immediately after mixing, virus-containing media was aspirated from each well and 1 mL plug media mixture was added. After plugs solidified, 500 µL D10 media (DMEM, 10% FBS, 8% sodium bicarbonate, 1x GlutaMAX, 1x MEM non-essential amino acids, 1% penicillin-streptomycin, 1 mM sodium pyruvate) was added to each well and plates were incubated overnight. Once syncytia were observed, media was aspirated and samples fixed for 10 minutes with 3.7% formaldehyde (Sigma 252549) diluted in 1x DPBS. Agar plugs were removed and 200 µL 1% crystal violet solution (Sigma V5265) was added to each well to stain cells for plaque counting and titer was determined according to the following equation:

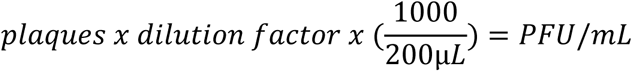

JHM^IA^ P1 stocks were passaged once in 17Cl-1 cells and virus titer for P2 Figure 5 was 1.1 x 10^6^ PFU/mL. To determine MOI, the following equation was used:

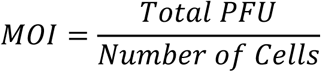

### acGFP-Sec61β stable FL83B cell generation

Following transfection of lentiviral plasmid DNA, lentivirus-containing media from 293T cells were collected at 24, 48 and 72 hours, pooled, and clarified for 10 minutes at 500 x g to remove cell debris. Virus was then concentrated using Lenti-X™ Concentrator reagent (Takara 631232). Briefly, 15 mL clarified virus supernatant was mixed with 5 mL Lenti-X Concentrator solution in a 50 mL conical and incubated at 4⁰C overnight. Sample was centrifuged at 1500 x g for 45 min at 4C, and supernatant was removed. Viral pellet was resuspended in 150 µL cold FL83B growth media, flash frozen with liquid nitrogen, and stored at −80°C before use. 280,000 FL83B cells were seeded per well in a 6-well plate for viral transduction the next day. Virus was rapidly thawed and diluted in 850 µL pre-warmed FL83B growth media supplemented with 10 µg/mL polybrene (Sigma TR-1003-G) before addition to cells. Mock infection controls (FL83B growth media + 10 µg/mL polybrene) were included to assess lentiviral toxicity. Virus and mock-infection media were removed after overnight incubation and replaced with 2 mL pre-warmed FL83B growth media. For maintenance and recovery, cells were split into large dishes to obtain a confluent 10-cm dish for cell sorting at CU Boulder’s Flow Cytometry Core Facility using a Miltenyi Biotec MACSQuant VYB Analyzer and 488 nm laser.

### Immunoblotting, immunostaining, and antibodies

For immunoblotting experiments, cell pellets were collected from a confluent 10 cm dish for each condition and subjected to lysis in 200 µL 1x XT Sample Buffer (BioRad #1610791) containing 1x XT reducing agent (BioRad #1610792) and 0.5 µL Benzonase® Nuclease (Millipore E1014) for 30 minutes on ice. Lysates were then clarified by centrifugation at 10,000 xg for 10 minutes at 4°C. Clarified lysates were loaded in duplicate in 12% Criterion™ XT Bis-Tris Protein Gels (BioRad #3450117) to probe for translocon subunits (Sec61α or EMC5) or mNeonGreen and run for 40 minutes at 200 V in either XT MOPS (BioRad 1610788, Sec61α) or MES (BioRad 1610789, EMC5) running buffer. Proteins were transferred onto Immun-Blot PVDF 0.2 µm Membranes (BioRad 1620177) for 1 hour at 75 V. Membranes were blocked in blocking buffer (5% nonfat milk solubilized in 1X TBST) for 1 hour followed by an overnight incubation at 4°C with primary antibodies diluted in blocking buffer. Primary antibodies rabbit anti-Sec61α (D7Q6V) recombinant monoclonal (Cell Signaling #14868) or rabbit anti-EMC5 (MMGT1) polyclonal (Bethyl laboratories #A305-833A). Membranes were incubated for 1 hour at room temperature with secondary antibodies goat anti-rabbit HPR conjugate (Sigma A6154) at 1:6000 diluted in blocking buffer. Membranes were visualized using SuperSignal West Pico PLUS Chemiluminescent Substrate (Thermo 34580).

For immunostaining of viral replication centers in infected cells (Figure 4D) and for imaging of HALO-Nsp6 transfected cells (Figure 4C), acGFP-Sec61β FL83B cells were seeded (as above) in 35-mm glass-bottom dishes the day before the experiment. Cells were transfected (as above) with 500 ng HALO-Nsp6 or infected with MHV JHM^IA^ at an MOI of 0.1. For infection, growth media was removed from the plate and replaced with virus diluted to 500 µL final in pre-warmed growth media. Cells were incubated with virus at 37°C and 5% CO_2_ for 1 hour with gentle rocking every 5 minutes. For the mock infection, 500 µL media without virus was substituted. The infectious media was then discarded and replaced with 2 mL fresh growth media. Infected and transfected cells were fixed at indicated hours post infection (mock, 8, 12, and 18 hpi) or transfection (24 hpt) with 2 mL pre-warmed fixation buffer (4% paraformaldehyde/EMS 15710, 1x FL83B growth media). Cells transfected with HALO-Nsp6 were incubated with 200 nM Janelia Fluor® JFX650 HaloTag® Ligand (Promega HT1070) for 15 min prior to fixation and were not further processed beyond permeabilization. Cells were fixed at 37⁰C for 10 minutes, washed 3 times with 2 mL room temperature 1x Dulbecco’s phosphate-buffered saline (DPBS/Gibco 14190250) and incubated with 2 mL room temperature permeabilization buffer (0.3% Triton X-100/ThermoFisher 28314, 1x DPBS) for 10 minutes. Fixed, infected cells were then incubated for 1 hour at room temperature with 200 µL blocking buffer (5% donkey serum/Sigma D9663, 0.1% Triton X-100, 1x DPBS) before immunostaining with primary mouse anti-dsRNA [J2] antibody (Sigma MABE1134) diluted 1:60 in blocking buffer and incubated overnight at 4°C on a tabletop rocker. Dishes were washed 3x with 2 mL room temperature 1x DPBS and incubated on a tabletop rocker for 1 hour at room temperature with 200 µL secondary donkey anti-mouse Alexa Fluor 568 (Thermo Fisher A10037) diluted 1:1000 in blocking buffer. Dishes were washed 3x with 2 mL room temperature 1x DPBS prior to imaging.

For TurboID proximity biotin ligase immunostaining experiments, FL83B cells transfected with TurboID-mNe-Nsp6, TurboID-mNe-empty vector, or mock transfected were treated as described above however, a Streptavidin Alexa Fluor 405 conjugated antibody (Invitrogen S32351) was used as the primary antibody at 1:100 (10 µg/mL) diluted in blocking buffer.

### Fluorescence microscopy

For fluorescence microscopy in Figure 1, Supplementary Figure S1 and immunostaining experiments (Figure 3 and 5), imaging was performed on a Zeiss Airyscan LSM 880 confocal line scanning microscope with Zeiss Zen 3.13 software using a Plan-Apochromat 63X (1.4 NA oil DIC M27) (Figure 1, 3, and S1) or 20X/0.8 objective in Airyscan fast (Figure S3) or LSM (Figure 5) mode. Dishes transfected with HALO-Nsp6 were incubated with 200 nM Janelia Fluor® JFX650 HaloTag® Ligand (Promega HT1070) for 15 min prior to imaging or fixation. Frame scanning was unidirectional scanning with no averaging, and images were taken in a single plane (64X, Figure 1, 3, 5, and S1) or with four z-planes which were processed to a maximum intensity projection (20X, 6 µm step size, S3). Pinhole sizes were >2.5 AU. The microscope is equipped with 405 nm (405-Streptavidin, BFP), 488 nm (mNeonGreen, acGFP), 561 nm (mCherry, Alexa Fluor 568), and 633 nm (JFX650) lasers with a beam splitter for the UV laser line. Emission filters used were BP 420-480 + BP 495-550, BP 495-550 + LP 570, BP 420-480 + LP 605, and BP 570-620 + LP 645. Detector gains were set to 700 V (405 nm) and 850 V (488, 561, and 633 nm) and digital gain 1.0. Scanning fields for Figure 1, 5 and S1 were 2X zoom, 1900 x 1900 pixels with pixel scaling of 0.04 x 0.04 µm, and 0.55 µs pixel dwell. For Figure 4, scanning field was 1.8X zoom, 1036 x 1036 with 0.07 x 0.07 µm pixel scaling, and 1.98 µs pixel dwell. For 20X objective images in Supplementary Figure 3 were 1X zoom, 1060 x 1060 pixels with pixel scaling of 0.40 x 0.40 x 2.00 µm, and 1.98 µs pixel dwell. All images taken on the 64X objective are 16-bit depth and 20X objective images are 8-bit depth.

Fluorescence microscopy for Mander’s colocalization coefficients and microscopy images in Figure 1 (Rtn3L, SigmaR1, and Climp63) and Supplementary Figure 2 were performed on live cells on a Yokogawa CSU-W1 SoRa spinning disk confocal using a 60X (1.42 NA) oil objective and 2.8X SoRa magnifiers. Camera used for acquisition is a Hamamatsu ORCA-Fusion Digital CMOS camera (C14440-20UP). Dishes transfected with HALO-Nsp6 were incubated with 200 nM Janelia Fluor® JFX650 HaloTag® Ligand (Promega HT1070) for 15 min prior to imaging. Imaging chambers were maintained at 37⁰C and 5% CO_2_ using Oko labs instrumentation. Laser excitation wavelengths used were 405 nm (BFP), 488 nm (mNeonGreen), 561 nm (mCherry), 640 nm (JFX650) with emission filters 455/50, 525/36, 605/52, and 705/72. SoRa (super-resolution) mode was utilized for image acquisition of a single frame and plane using Nikon NIS-Elements software. Laser power and exposure times were as follows: 405 nm, 80%, 200 ms; 488 nm, 80%, 300 ms; 561 nm, 80%, 200 ms, 640 nm 20%, 200 ms. Image dimensions were 2304 x 2304 pixels with 38.69 nm/pixel effective super-resolution and were not deconvolved.

### FRAP and FLIP acquisition and analysis

For fluorescence recovery after photobleaching (FRAP) and fluorescence loss in photobleaching (FLIP) experiments, cells were incubated with 200 nM Janelia Fluor® JFX650 HaloTag® Ligand (Promega HT1070) for 15 min prior to imaging. Imaging was performed at 37⁰C on a Zeiss Airyscan LSM 880 confocal line scanning microscope with Zeiss Zen 3.13 software using a Plan-Apochromat 63X (1.4 NA oil DIC M27) objective in LSM mode. Scanning was unidirectional frame scanning with no averaging, and a single plane. Detector gains were set to 850 V (488 nm) and 700 V (633 nm) and digital gain 1.0. Pinhole size was 2.53 AU. All images were 16-bit depth. 488 nm laser with BP 420-480 + BP 495-550 emission filters was used to acquire mNeonGreen (Sec61α and SRPRβ) and 633 nm laser with BP 570-620 + LP 645 emission filters was used to acquire HALO (Nsp6). An image of the HALO-Nsp6 channel image was taken once at the beginning of acquisition. Scanning fields were 7X zoom, 164 x 164 pixels with 0.10 µm x 0.10 µm pixel scaling, and 0.92 µs pixel dwell (FRAP) or 2.5X zoom,1552 x 1552 pixels with 0.03 µm x 0.03 µm pixel scaling, and 0.80 µs pixel dwell (FLIP).

ROIs were defined using Zeiss Zen Blue 3.13 imaging software prior to image acquisition and the Bleaching tool was used for bleaching setup. 2-3 µm (FRAP) regions of interest (ROIs) or 10-15 µm (FLIP) ROIs were bleached (t = 0) down to ∼10% signal intensity using high intensity (80%) pulses with the 488 nm laser. For FRAP and FLIP time-course, laser power for the 488 nm laser was set to 3%. 20 (FRAP) or 10 (FLIP). Pre-bleach frames were acquired before bleaching for baseline normalizations. ROIs were re-bleached every 50 seconds for FLIP experiments. For FRAP experiments, 74 second time-lapses collected at 40 ms intervals were acquired to measure fluorescence recovery. For FLIP experiments, time lapses were acquired for 945 seconds at 5 sec intervals to observe ∼90% loss of fluorescence signal in the controls. ROIs were drawn using the Zeiss Zen Blue 3.13 software prior to image acquisition and ROI intensity data was exported from the Zen Black software. Pre-bleach frame mean intensity (T_prebleach_) was used for recovery curve normalizations. To generate normalized recovery/decay for curve fitting, a double normalization of recovery/decay signal was performed using the equation below which has been published previously^87^, where T_t_ is the mean intensity (at time t) within the total imaging region, I_t_ is the mean intensity (at time t) within the bleached ROI, background (BG) was determined by the mean intensity in regions devoid of ER signal and away from the bleached ROI. For FLIP normalizations, T_t_ is instead the mean intensity (at time t) of a neighboring cell within the same imaging region:

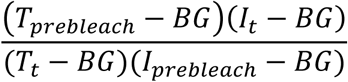

To extract the half-time recovery (t_1/2_), recovery rate (k), and mobile fractions, FRAP and FLIP data was fit to a nonlinear least squares regression for a single exponential using GraphPad Prism version 10.6.1 one phase association (FRAP):

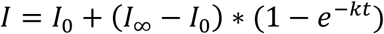

Or one phase decay (FLIP):

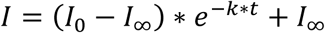

Where k is the recovery/decay rate determined by best fit, I_0_ = the mean intensity at time (t) 0 and I_∞_ = the mean maximum intensity recovered/lost in the imaging time course. Half-time recovery/decay (t_1/2_) = ln(2)/k, % mobile fraction = (I_∞_-I_0_)*100.

### qPCR sample preparation and RNA isolation

FL83B cells were seeded (220,000 cells/well) in a 6-well plate the day before to achieve ∼70% confluence on the day of transfection/infection. Cells were transfected (as above) with 500 ng HALO-Nsp6 or infected with MHV JHM^IA^ at an MOI of 0.1. For infection, growth media was removed from the well and replaced with virus diluted to 500 µL final in pre-warmed growth media. Cells were incubated with virus at 37⁰C and 5% CO_2_ for 45 min with gentle rocking every 5 minutes. For the mock infection, 500 µL media without virus was substituted. The infectious media was then discarded and replaced with 2 mL fresh pre-warmed growth media. For timepoint 0, virus was collected immediately after the 45 min virus incubation after washing 2x with ice-cold 1x DPBS and cell pellets were flash frozen in liquid nitrogen. Infected cells were collected at the indicated time points (3 hpi, 8 hpi, 12 hpi, 18 hpi, and 24 hpi) and transfected cells were collected 24 hours post transfection. In both cases, cells were washed 2x with ice-cold 1x DPBS prior to collection and cell pellets were flash frozen in liquid nitrogen. Virus and host RNA was isolated from cell pellets using the Monarch Total RNA Miniprep Kit (NEB # T2010S) according to the manufacturer’s included protocol for cultured mammalian cells. RNA was eluted in 50 µl of nuclease-free water and the concentration was determined using a Nanodrop 2000 spectrophotometer (Thermo Fisher Scientific).

### RT-qPCR

First-strand cDNA synthesis was performed on all RNA samples using the LunaScript® RT Master Mix Kit (NEB #E3025L) combined with the Random Primer Mix (NEB #S1330S) according to the manufacturer’s included protocol. For each biological replicate, all first strand cDNA synthesis reactions were performed using an equal weight of template RNA (500 ng). Subsequently, all cDNA reactions were diluted 3-fold in nuclease-free water, from a 20 µl volume to a final 60 µl.

Quantitative PCR reactions were performed using the Luna Universal qPCR Master Mix (NEB #M3003L) according to the manufacturer’s protocol. For each biological replicate, two technical replicate reactions were run on an Applied Biosystems Fast 7500 instrument according to the fast protocol. Threshold cycle (Ct), which is the number of cycles when a target amplicon is detected above threshold, values from all samples were normalized to a housekeeping gene before relative quantitation. All samples isolated from whole cells were normalized to beta-actin RNA. Relative quantitation was performed according to the delta Ct method (ddCt). Delta Ct (*dCt*), which represents normalization to the housekeeping gene is calculated using the following equation:

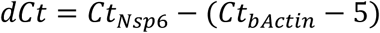

where *Ct*Nsp6 is the mean threshold cycle of technical replicates for Nsp6 RNA in a sample, *Ct*_bActin_ is the mean threshold cycle of technical replicates for the housekeeping gene in the same sample. A constant of 5 was subtracted from *Ct*_bActin_ values to avoid negative numbers. Ct values represent the cycle at which the target nucleic acid was detected, meaning lower values indicate higher amounts in the sample on a logarithmic scale.

### TurboID proximity labeling

For proximity labeling biotin ligase experiments using TurboID, FL83B cells were transfected as described above in three 10-cm dishes for each condition (TurboID-mNe-Nsp6, TurboID-mNe-empty vectors, mock transfection control) to obtain ∼ 4 mg of total protein each. Each dish was treated with 500 µM biotin (Sigma B4501) for 1 hour, washed 3x with ice-cold 1x DPBS, and scraped in 1 mL lysis buffer (25 mM Tris-HCl pH 7.4, 200 mM NaCl, 1mM EDTA, 1% NP-40, 1% SDS, 5% β-mercaptoethanol, 1x Halt™ Protease Inhibitor Cocktail/Thermo Fisher 87786). Cells were incubated on ice for 30 minutes and vortexed briefly every 5 minutes to complete lysis. Cell debris were clarified at 10,000 x g for 10 minutes at 4°C. To remove interfering agents in the lysis buffer, proteins were precipitated in acetone by mixing 1 mL sample with 4 mL ice-cold 100% acetone, vortexed, and incubated at −20°C for 90 min. Replicates were pooled and total protein was pelleted at 21,000 x g for 10 min. Supernatants were decanted and pellets dried at room temperature for 20 min and flash frozen with liquid nitrogen. Protein pellets were sent on dry ice to Sanford Burnham Prebys Medical Discovery Institute proteomics facility for processing.

### Mass Spectrometry and data analysis

The following steps were provided by and performed at Sanford Burnham Prebys. The protein pellet was dissolved in 8M urea, 50 mM ammonium bicarbonate (ABC) and the solution was centrifuged at 14,000 x g for 15 minutes to remove insoluble particles. Supernatant protein concentration was determined using a bicinchoninic acid (BCA) protein assay (Thermo Scientific). Disulfide bridges were reduced with 5 mM tris(2-carboxyethyl)phosphine (TCEP) at 30°C for 60 min, and cysteines were subsequently alkylated with 15 mM iodoacetamide (IAA) in the dark at room temperature for 30 min. Affinity purification was carried out in a Bravo AssayMap platform (Agilent) using AssayMap streptavidin cartridges (Agilent). Briefly, cartridges were first primed with 50 mM ammonium bicarbonate, and then proteins were slowly loaded onto the streptavidin cartridge. Background contamination was removed with 8M urea, 50 mM ammonium bicarbonate. Finally, cartridges were washed with Rapid digestion buffer (Promega, Rapid digestion buffer kit) and proteins were subjected to on-cartridge digestion with mass spec grade Trypsin/Lys-C Rapid digestion enzyme (Promega, Madison, WI) at 70C for 1h. Digested peptides were then desalted in the Bravo platform using AssayMap C18 cartridges, and dried down in a SpeedVac concentrator.

Prior to LC-MS/MS analysis, dried peptides were reconstituted with 2% ACN, 0.1% FA and concentration was determined using a NanoDropTM spectrophometer (ThermoFisher). Samples were then analyzed by LC-MS/MS using a Proxeon EASY-nanoLC system (ThermoFisher) coupled to a Orbitrap Fusion Lumos Tribid mass spectrometer (Thermo Fisher Scientific). Peptides were separated using an analytical C18 Aurora column (75µm x 250 mm, 1.6 µm particles; IonOpticks) at a flow rate of 300 nL/min (60C) using a 75-min gradient: 2% to 6% B in 1 min, 6% to 23% B in 45 min, 23% to 34% B in 28 min, and 34% to 48% B in 1 min (A= FA 0.1%; B=80% ACN: 0.1% FA). The mass spectrometer was operated in positive data-dependent acquisition mode. MS1 spectra were measured in the Orbitrap in a mass-to-charge (m/z) of 375 – 1500 with a resolution of 60,000. Automatic gain control target was set to 4 x 10^5 with a maximum injection time of 50 ms. The instrument was set to run in top speed mode with 1-second cycles for the survey and the MS/MS scans. After a survey scan, the most abundant precursors (with charge state between +2 and +7) were isolated in the quadrupole with an isolation window of 0.7 m/z and fragmented with HCD at 30% normalized collision energy. Fragmented precursors were detected in the ion trap as rapid scan mode with automatic gain control target set to 1 x 10^4^ and a maximum injection time set at 35 ms. The dynamic exclusion was set to 20 seconds with a 10 ppm mass tolerance around the precursor.

All mass spectra from were analyzed with MaxQuant software version 1.6.11.0. MS/MS spectra were searched against the Mus musculus Uniprot protein sequence database (downloaded in Mar 2022), Nsp6 and control constructs, and GPM cRAP sequences (commonly known protein contaminants). Precursor mass tolerance was set to 20ppm and 4.5ppm for the first search where initial mass recalibration was completed and for the main search, respectively. Product ions were searched with a mass tolerance 0.5 Da. The maximum precursor ion charge state used for searching was 7. Carbamidomethylation of cysteine was searched as a fixed modification, while oxidation of methionine and acetylation of protein N-terminal were searched as variable modifications. Enzyme was set to trypsin in a specific mode and a maximum of two missed cleavages was allowed for searching. The target-decoy-based false discovery rate (FDR) filter for spectrum and protein identification was set to 1%.

### Software, image processing, and statistical methods

For Mander’s colocalization analysis^32^ between translocons, one 10 x 10 µm inset was cropped from the periphery of each cell and the mean background was subtracted for each channel using math processing in FIJI v1.54f^88^. Otsu’s method was applied to each channel to determine thresholding values using the FIJI Auto Threshold plugin^89^. To score Mander’s colocalization, thresholding values from Otsu’s method were inputted for each channel in the JACoP FIJI plugin. To simulate random chance, the KDEL and Nsp6 channels were rotated 90° to the right and the same parameters were applied for Mander’s colocalization scoring. Cartoons were drawn, figures designed and compiled using Adobe Illustrator 2025. All processing of microscopy images for image presentation and linear adjustments to correct for photobleaching in FRAP and FLIP examples were done in FIJI. MCC, FRAP, FLIP, and proteomics datasets were graphed using GraphPad Prism 10.6.1. All datasets are represented by three biological replicates. FRAP and FLIP replicates each contain n = 5 and MCC replicates contain n = 15 cells. Mean, standard error of the mean, and standard deviation were calculated using GraphPad Prism 10.6.1. P-values were calculated by performing an unpaired t test from the mean and standard error of three biological replicates using GraphPad Prism 10.6.1. A threshold of P < 0.05 was used to determine significance. For western blot quantitation, band intensities for endogenous protein and exogenous from two western blots were measured using the FIJI western blot analyzer plugin and plotted using GraphPad Prism 10.6.1.

To identify top ER proteomic hits enriched in the Nsp6 samples, TurboID-mNe-Nsp6 iBAQ values for each proteomic hit were divided by the iBAQ value from the cytoplasmic (TurboID-mNe empty vector) or the non-specific bead-binding (mock transfection) controls, choosing the iBAQ control value that was greatest. Many proteomic hits were not present in the control samples (iBAQ = 0), therefore, a constant of 1.0 was added to all iBAQ values to perform the division. For determining localization of proteomic hits (i.e. ER proteins), an in-house Python script was used to query the Uniprot REST API and extract subcellular localization data using protein Accession IDs. iBAQ values for ER proteins were plotted on a Log(2) scale and translocon machinery and translocon-associated proteins were identified in the list of top hits and labeled manually on the plot.

