## Supplemental Material for "β-Coronavirus Nsp6 hijacks host ER translocation machineries into viral replication centers"

**SUPPLEMENTARY MATERIAL**

Figure S1. ***MHV nonstructural protein Nsp6 remodels ER domains and recruits Sec61 in Cos-7 cells.*** (A,B) Representative FL83B cells transfected with BFP-Sec61β (blue) co-transfected with (A) mNe-Nsp3 (green) and HALO-Nsp6 (magenta), or (B) mCh-Nsp4 (green) and HALO-Nsp6 (magenta). (C-F) Representative images of COS-7 cells co-transfected with (C) mNe-Nsp3, mCh-Nsp4, HALO-Nsp6 (conjugated to JFX 650) and with either BFP-KDEL or (D) BFP-Sec61β. (E) mNe-Nsp3 (top), mCh-Nsp4 (middle), or HALO-Nsp6 (bottom) (conjugated to JFX 650) and with either BFP-KDEL (left) or (F) BFP-Sec61β (right). (A-F) Inset white boxes shown in left panels are magnified in panels on the right. Scale bars = 5 μm.

Figure S2. **Translocon associated components can be sequestered by Nsp6.** (A,C) Immuno-blot example of (A) Sec61α or (B) EMC5 in FL83B cells mock transfected (lane 1), transfected withSec61α-mNe, or EMC5-mNe (lane 2), or with Sec61α-mNe or mNe-EMC5 and HALO-Nsp6 (lane 3). Molecular weight standards are shown. Black arrow heads point to endogenous and exogenous expression bands for Sec61α (left) or EMC5 (right). Relative band intensity of transiently expressed Sec61α or EMC5 was divided by the endogenous band intensity and plotted. For Sec61α-mNe, the double band was included in the intensity ratio. n = 2 western blots. (B, D-I) Representative images of FL83B expressing HALO-Nsp6 (magenta) with BFP-KDEL (green) and (B) mNe-EMC4, (D) GFP-Ergic53, (E) mNe-Ssr3, (F)

Ssr1-mNe, (G) Spcs1-mNe, (H) Spcs2-mNe, or (I) mNe-Spcs3. Individual channels are shown next to the merged inset with labels. Inset white boxes shown in left panels are magnified in panels on the right. Scale bars = 5 µm.

Figure S3. **acGFP-Sec61β FL83B cells infected with MHV show Sec61β signal overlapping dsRNA.** (A) Lower magnification (20X) images of 0.1 MOI MHV JHM$^{IA}$ infected FL83B cells at 0 and 8 hpi. acGFP-Sec61β is shown in green and replication centers are shown in magenta (immune-stained with an antibody against dsRNA). Scale bars = 50 µm.

Table S1. **Biotinylated ER proteins from mNe-TurboID-Nsp6 experiments.** Rank, Protein IDs, and Gene names for the top 30 *mus musculus* ER proteins identified by mass spectrometry. See methods for detailed explanation of protein rank determination.

**Supplementary Table 1**

| Rank | Protein IDs | Protein names | Gene names |
|---|---|---|---|
| 1 | | Nsp6 | Nsp6 |
| 2 | Q9DCF9 | Translocon-associated protein subunit gamma | Ssr3 |
| 3 | Q9D958 | Signal peptidase complex subunit 1 | Spcs1 |
| 4 | Q6ZWQ7 | Signal peptidase complex subunit 3 | Spcs3 |
| 5 | Q9CRA4 | Methylsterol monooxygenase 1 | Msmo1 |
| 6 | P35821 | Tyrosine-protein phosphatase non-receptor type 1 | Ptpn1 |
| 7 | P70245 | 3-beta-hydroxysteroid-Delta(8),Delta(7)-isomerase | Ebp |
| 8 | Q9CZX9 | ER membrane protein complex subunit 4 | Emc4 |
| 9 | Q921J2 | GTP-binding protein Rheb | Rheb |
| 10 | Q80TA1 | Ethanolaminephosphotransferase 1 | Ept1 |
| 11 | Q9ERY9 | Probable ergosterol biosynthetic protein 28 | ORF11 |
| 12 | Q5RL79 | Keratinocyte-associated protein 2 | Krtcap2 |
| 13 | Q8K4X7 | 1-acyl-sn-glycerol-3-phosphate acyltransferase delta | Agpat4 |
| 14 | O35166 | Golgi SNAP receptor complex member 2 | Gosr2 |
| 15 | O88455 | 7-dehydrocholesterol reductase | Dhcr7 |
| 16 | Q9CZW4 | Long-chain-fatty-acid--CoA ligase 3 | Acsl3 |
| 17 | P45878 | Peptidyl-prolyl cis-trans isomerase FKBP2 | Fkbp2 |

| | | | |
|---|---|---|---|
| 18 | Q80TN4 | DnaJ homolog subfamily C member 16 | Dnajc16 |
| 19 | Q80W54 | CAAX prenyl protease 1 homolog | Zmpste24 |
| 20 | Q9CQT9 | Uncharacterized protein C20orf24 homolog | OPTI |
| 21 | Q9CQZ0;Q921I0 | ORM1-like protein 2;ORM1-like protein 1 | Ormdl2;Ormdl1 |
| 22 | Q9D2C7 | Bax inhibitor 1 | Tmbim6 |
| 23 | Q6PA06;Q8BH66 | Atlastin-2 | Atl2 |
| 24 | Q99KK1 | Receptor expression-enhancing protein 3 | Reep3 |
| 25 | O09005 | Sphingolipid delta(4)-desaturase DES1 | Degs1 |
| 26 | Q8VE91 | Protein FAM134B | Fam134b |
| 27 | O35623 | BET1 homolog | Bet1 |

Figure S1

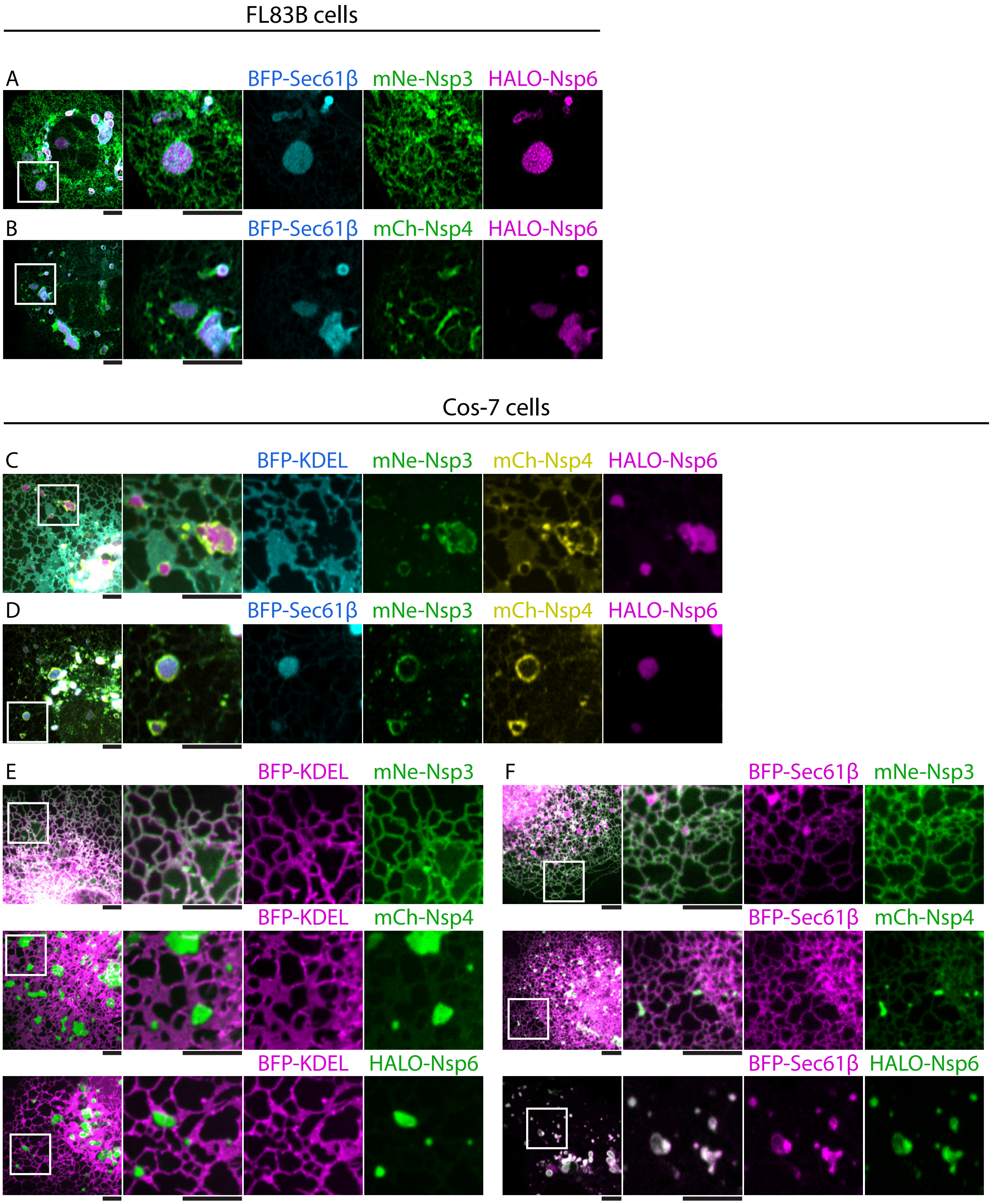

FL83B cells
A
BFP-Sec61β
mNe-Nsp3
HALO-Nsp6
B
BFP-Sec61β
mCh-Nsp4
HALO-Nsp6
Cos-7 cells
C
BFP-KDEL
mNe-Nsp3
mCh-Nsp4
HALO-Nsp6
D
BFP-Sec61β
mNe-Nsp3
mCh-Nsp4
HALO-Nsp6
E
BFP-KDEL
mNe-Nsp3
BFP-KDEL
mCh-Nsp4
BFP-KDEL
HALO-Nsp6
F
BFP-Sec61β
mNe-Nsp3
BFP-Sec61β
mCh-Nsp4
BFP-Sec61β
HALO-Nsp6

Figure S2

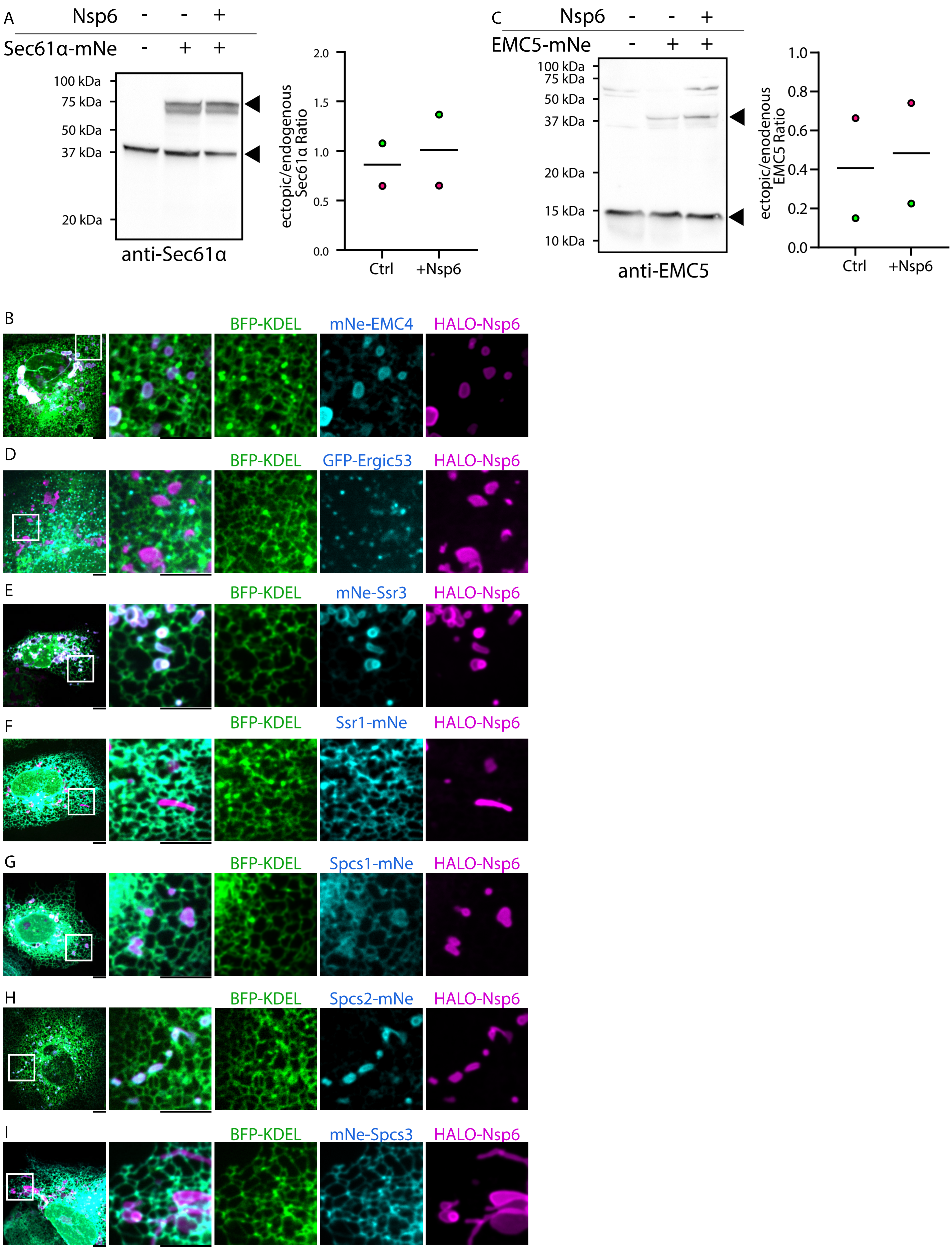

A
Nsp6 - - +
Sec61α-mNe - + +
100 kDa
75 kDa
50 kDa
37 kDa
20 kDa
anti-Sec61α
ectopic/endogenous Sec61α Ratio
2.0
1.5
1.0
0.5
0.0
Ctrl
+Nsp6
C
Nsp6 - - +
EMC5-mNe - + +
100 kDa
75 kDa
50 kDa
37 kDa
20 kDa
15 kDa
10 kDa
anti-EMC5
ectopic/enodenous EMC5 Ratio
1.0
0.8
0.6
0.4
0.2
0.0
Ctrl
+Nsp6
B
BFP-KDEL
mNe-EMC4
HALO-Nsp6
D
BFP-KDEL
GFP-Ergic53
HALO-Nsp6
E
BFP-KDEL
mNe-Ssr3
HALO-Nsp6
F
BFP-KDEL
Ssr1-mNe
HALO-Nsp6
G
BFP-KDEL
Spcs1-mNe
HALO-Nsp6
H
BFP-KDEL
Spcs2-mNe
HALO-Nsp6
I
BFP-KDEL
mNe-Spcs3
HALO-Nsp6

Figure S3

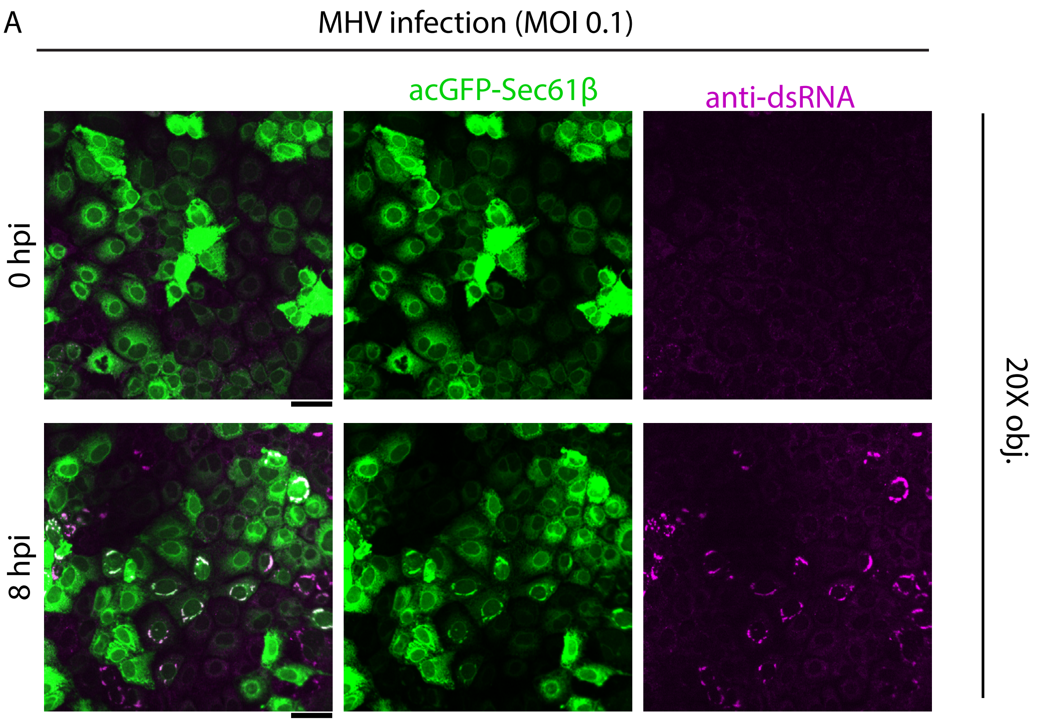
A
MHV infection (MOI 0.1)
acGFP-Sec61β
anti-dsRNA
0 hpi
8 hpi
20X obj.
